# Small DNA elements that act as both insulators and silencers in plants

**DOI:** 10.1101/2024.09.13.612883

**Authors:** Tobias Jores, Nicholas A. Mueth, Jackson Tonnies, Si Nian Char, Bo Liu, Valentina Grillo-Alvarado, Shane Abbitt, Ajith Anand, Stéphane Deschamps, Scott Diehn, Bill Gordon-Kamm, Shuping Jiao, Kathy Munkvold, Heather Snowgren, Nagesh Sardesai, Stanley Fields, Bing Yang, Josh T. Cuperus, Christine Queitsch

## Abstract

Insulators are *cis*-regulatory elements that separate transcriptional units, whereas silencers are elements that repress transcription regardless of their position. In plants, these elements remain largely uncharacterized. Here, we use the massively parallel reporter assay Plant STARR-seq with short fragments of eight large insulators to identify more than 100 fragments that block enhancer activity. The short fragments can be combined to generate more powerful insulators that abolish the capacity of the strong viral 35S enhancer to activate the 35S minimal promoter. Unexpectedly, when tested upstream of weak enhancers, these fragments act as silencers and repress transcription. Thus, these elements are capable of both insulating or repressing transcription dependent upon regulatory context. We validate our findings in stable transgenic *Arabidopsis*, maize, and rice plants. The short elements identified here should be useful building blocks for plant biotechnology efforts.

## Main text

Precise control of gene expression is crucial for plants to grow and develop in a changing environment. Genomic approaches to study plant gene regulation have focused mainly on promoters and enhancers^1–7^. In contrast, repressive elements such as silencers and insulators have received far less attention. Insulators compartmentalize genomes into discrete transcriptional units^8^. Insulators have one or both of two principal functions^9^: They block enhancers from interacting with core promoters (enhancer-blocking insulators), or they form barriers against the spread of repressive heterochromatin (barrier insulators). Enhancer-blocking insulators are defined by their ability to act when situated between an enhancer and promoter, but not when the order is reversed such that the enhancer is closer to the promoter than the insulator^8^. Insulators are thought to prevent ectopic gene expression, maintain chromatin accessibility, and enable differentially regulated genes to reside in close proximity to one another^10^.

To date, most research on insulators has been performed in animal models^11^. In contrast, only a handful of plant sequences have been shown to act as insulators in transient or stable transgenic plant reporter assays^12,13^. For example, the Transformation Booster Sequence (TBS) from *Petunia hybrida,* the β-phaseolin gene from *Phaseolis vulgaris*, and a *gypsy*-like sequence from *Arabidopsis thaliana* function as enhancer-blocking insulators in transgenic plants^14–16^. In addition, a few heterologous sequences show enhancer-blocking insulator activity in plants, including λ-EXOB from phage λ, BEAD-1C from humans, and UASrpg from yeast^17,18^. In these studies, insulator activity was inferred from β-glucuronidase (GUS) staining or fluorescence of a reporter gene, both measures with limitations of dynamic range, quantification accuracy, and throughput.

Because the enhancer-blocking activity of insulators is detected as reduced transcription in the commonly used reporter assays, care must be taken to distinguish between insulators and silencers, which could also cause reduced transcription in these assays. Silencers recruit repressive transcription factors and, like enhancers, can act in a position-independent manner (*i.e.* upstream or downstream of an enhancer)^6,19–22^. This position-independency is thought to be a key difference between silencers and insulators that differentiates between the two element types.

To date, no general principles are known that typify insulator or silencer function in plants nor are there high-throughput methods to identify these elements. Short and strong insulators will facilitate synthetic biology applications to ensure predictable expression of transgenes, blocking inappropriate enhancer-promoter interactions and alleviating chromatin position effects^12^. Similarly, silencers will enable fine-tuning of transgene expression and minimize expression noise. Furthermore, understanding the sequence features of functional plant insulators and silencers will allow targeting similar elements in plant genomes to engineer gene expression.

Here, we applied Plant STARR-seq, a massively parallel reporter assay, to test the insulator and silencer activity of over 100 short (170 bp) fragments derived from either previously described enhancer-blocking insulators or two novel synthetic insulator sequences. Our assay distinguishes enhancer-blocking activity from transcriptional repression and reveals that the insulator-derived elements harbor both insulator-like and silencer-like activities. Promising elements were tested and verified in stable transgenic *Arabidopsis*, rice, and maize plants.

## Results

### Plant STARR-seq detects the activity of enhancer-blocking insulators

Plant STARR-seq can identify and characterize *cis*-regulatory elements^3,4,7,23,24^. To test whether Plant STARR-seq can identify enhancer-blocking insulators, we created a reporter construct consisting of a barcoded green fluorescent protein (GFP) gene under the control of a 35S minimal promoter coupled to a 35S enhancer; insulator candidates were placed between this enhancer and promoter (Extended Data Fig. 1a). We selected four heterologous sequences that show insulator activity in plants (λ-EXOB, BEAD-1C, UASrpg, and a *Drosophila* gypsy element, refs. ^12,17,18,25^), and two synthetic sequences (sIns1 and sIns2) for which preliminary data suggested they might act as insulators. The synthetic sequences sIns1 and sIns2 derive from a plasmid backbone and a human codon-optimized coding sequence of Cas9, respectively. The insulator candidate sequences were cloned in the forward or reverse orientation, and their insulator activity was determined by Plant STARR-seq in tobacco (*Nicotiana benthamiana*) leaves and maize (*Zea mays*) protoplasts. Constructs without the 35S enhancer and without an insulator (noEnh); with the 35S enhancer and without an insulator (noIns) were included as controls. We measured insulator activity as reduced enrichment compared to the enrichment of the no insulator (noIns) control. Except for the gypsy element, the other five tested insulator candidates resulted in reduced enrichment, indicating that they function as enhancer-blocking insulators in this assay (Extended Data Fig. 1b). The gypsy element shows enhancer-blocking and barrier insulator activities in *Drosophila*^26^; however, it lacks enhancer-blocking activity in plants^25^, consistent with our results (Extended Data Fig. 1b). For some of the insulators, we observed orientation-dependent activity (Extended Data Fig. 1b). Taken together, we demonstrate that Plant STARR-seq reproducibly (Extended Data Fig. 2) measures insulator activity.

The large size of known enhancer-blocking insulators precludes their application in plant biotechnology^12^. To identify short sequences with insulator activity, we array-synthesized overlapping 170-bp fragments of each of the six insulators in addition to two plant sequences with insulator activity (β-phaseolin and TBS), and measured the enhancer-blocking activity of these fragments (Fig. 1a). Many fragments retained partial insulator activity in tobacco and maize (Fig. 1b and Supplementary Data 1), but their activity varied between the two assay systems, pointing to species-specific differences (Fig. 1c).

**Fig. 1.**
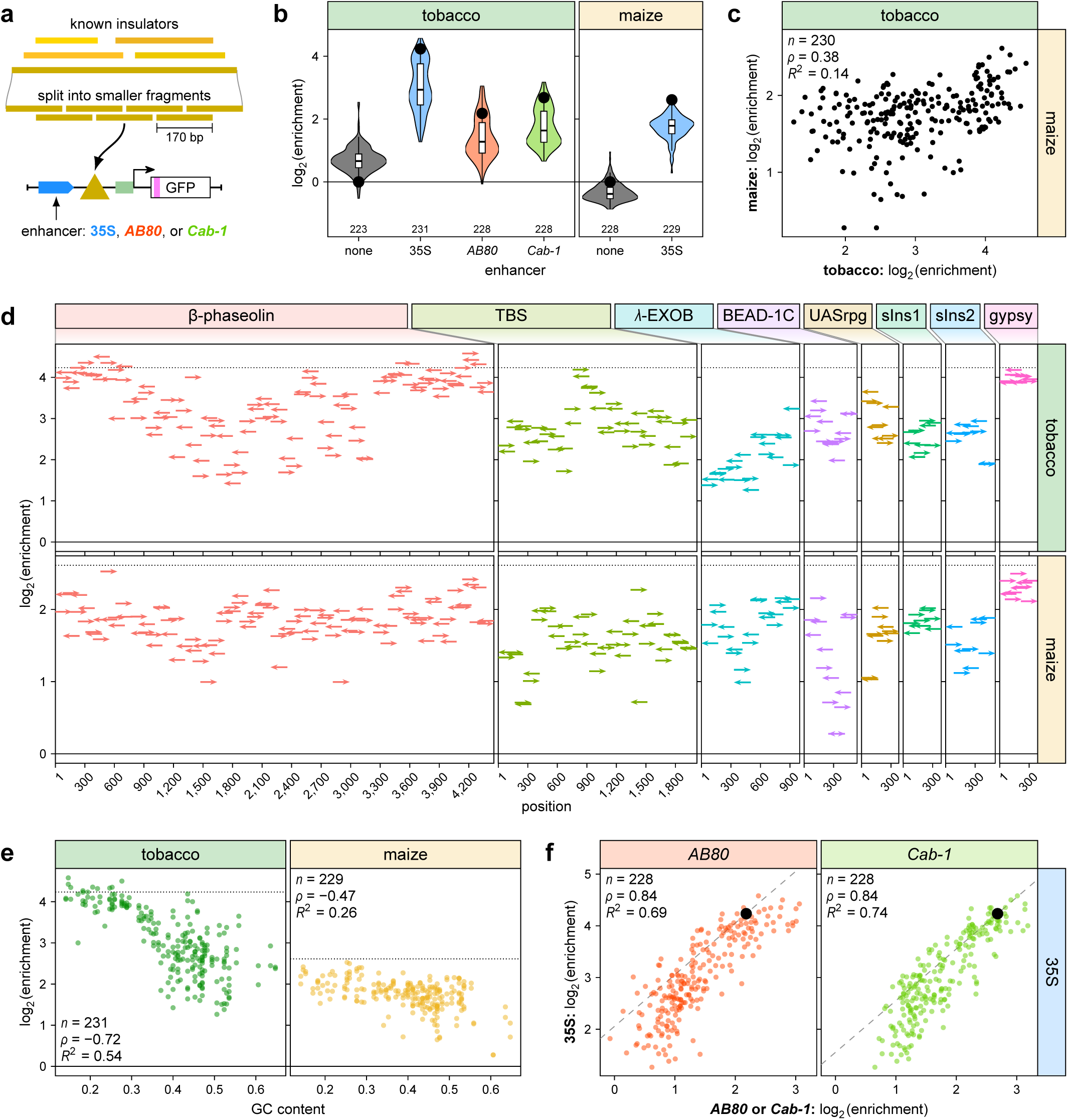
Short fragments exhibit enhancer-blocking insulator activity. **a**, Known insulators were split into partially overlapping 170-bp fragments. The insulator fragments were cloned in the forward or reverse orientation between a 35S, *AB80*, or *Cab-1* enhancer and a 35S minimal promoter driving the expression of a barcoded GFP reporter gene. Constructs without an enhancer (none) but with insulator fragments were also created. **b**, All insulator fragment constructs were pooled and subjected to Plant STARR-seq in tobacco leaves (tobacco) and maize protoplasts (maize). Reporter mRNA enrichment was normalized to a control construct without an enhancer or insulator (noEnh; log2 set to 0). The enrichment of a control construct without an insulator is indicated as a black dot. Violin plots represent the kernel density distribution and the box plots inside represent the median (center line), upper and lower quartiles, and 1.5× interquartile range (whiskers) for all corresponding constructs. Numbers at the bottom of each violin indicate the number of samples in each group. **c**, Correlation between the enrichment of insulator fragments in constructs with the 35S enhancer in tobacco leaves and maize protoplasts. **d**, Enrichment of constructs with insulator fragments cloned between the 35S enhancer and minimal promoter. The position along the full-length insulator and the orientation (arrow pointing right, fwd; arrow pointing left, rev) of the fragments is indicated by arrows. **e**, Correlation between insulator fragment enrichment and GC content for constructs with the 35S enhancer. **f**, Correlation between insulator fragment enrichment in tobacco leaves in constructs with the indicated enhancers. The dashed line represents a y = x line fitted through the point corresponding to a control construct without an insulator (black dot). Pearson’s *R*^2^, Spearman’s *ρ*, and number (*n*) of constructs are indicated in **c**, **e**, and **f**.

Overall, seven of the eight insulators, excluding only the gypsy element, harbored clusters of fragments that partially blocked the 35S enhancer (Fig 1d). This clustering of active fragments is likely driven by local nucleotide composition because GC content strongly correlated with a fragment’s insulator activity (Fig. 1e). However, GC content does not fully explain insulator activity: Many insulator-derived fragments showed orientation-dependent activity (Fig. 1d). Furthermore, we tested the insulator-derived fragments with the *AB80* enhancer from *Pisum sativum* and *Cab-1* enhancer from *Triticum aestivum*, which drive the expression of chlorophyl a-b binding proteins, and found that the activity of these fragments was largely enhancer-independent (Fig. 1b,f).

To validate our findings, we measured insulator activity in stable transgenic plants (Fig. 2). Full-length insulators and fragments thereof showed enhancer-blocking insulator activity in *Arabidopsis*, rice (*Oryza sativa*), and maize, well correlated with the Plant STARR-seq results (Fig. 2 and Extended Data Fig. 3). In maize, we measured insulator activity in four tissues (leaf, stalk, silk, and husk) and two developmental stages (V6 and R1) and obtained similar results, indicating that these insulators do not act in a tissue-specific manner (Fig. 2k).

**Fig. 2.**
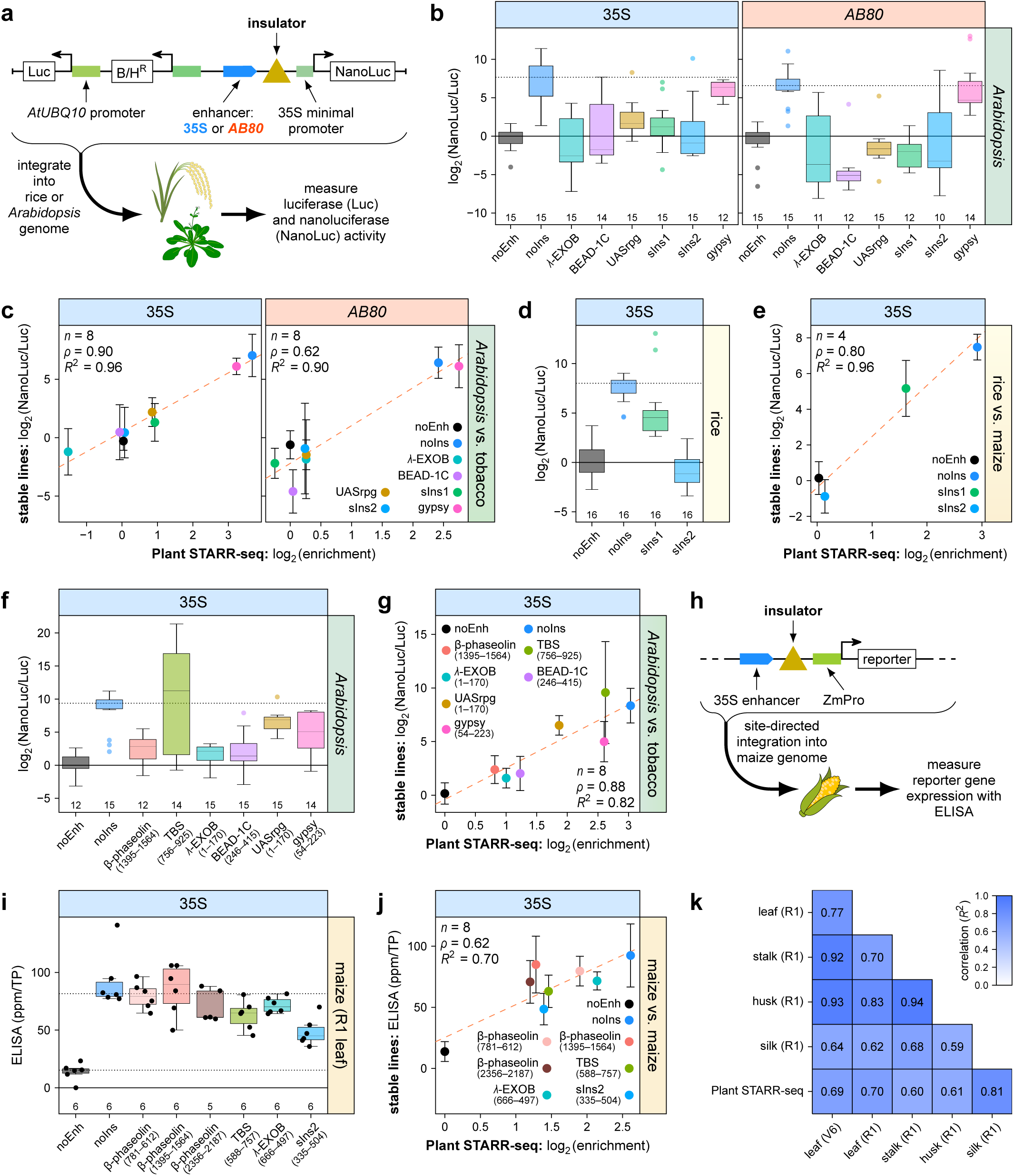
Insulators are active in stable transgenic lines in *Arabidopsis*, rice, and maize. **a**, Transgenic *Arabidopsis* and rice lines were generated with T-DNAs harboring a constitutively expressed luciferase (Luc) gene and a nanoluciferase (NanoLuc) gene under control of a 35S minimal promoter coupled to the 35S or *AB80* enhancer (as indicated above the plots) with insulator candidates inserted between the enhancer and promoter. Nanoluciferase activity was measured in at least 4 plants from these lines and normalized to the activity of luciferase. The NanoLuc/Luc ratio was normalized to a control construct without an enhancer or insulator (noEnh; log2 set to 0). **b**,**c**, The activity of full-length insulators was measured in *Arabidopsis* lines (**b**) and compared to the corresponding results from Plant STARR-seq in tobacco leaves (**c**). **d**,**e**, The activity of synthetic full-length insulators was measured in rice lines (**d**) and compared to the corresponding results from Plant STARR-seq in maize protoplasts (**e**). **f**,**g**, The activity of insulator fragments was measured in *Arabidopsis* lines (**f**) and compared to the corresponding results from Plant STARR-seq in tobacco leaves (**g**). **h**, For transgenic maize lines, a reporter gene driven by a moderate-strength constitutive promoter (ZmPro) and an upstream 35S enhancer was created and insulator fragments were inserted between the enhancer and promoter. The reporter gene cassette was inserted in the maize genome by site-directed integration and the expression of the reporter gene was measured in various tissues/developmental stages by ELISA. **i**,**j**, The activity of insulator fragments was measured in R1 leaves of transgenic maize lines (**i**) and compared to the corresponding results from Plant STARR-seq in maize protoplasts (**j**). **k**, Correlation (Pearson’s *R*^2^) between the expression of all tested constructs across different tissues and developmental stages. The correlation with Plant STARR-seq results from maize protoplasts is also shown. Box plots in **b**, **d**, (**f**), and (**i**) represent the median (center line), upper and lower quartiles (box limits), 1.5× interquartile range (whiskers), and outliers (points) for all corresponding samples from two to three independent replicates. Numbers at the bottom of each box plot indicate the number of samples in each group. The enrichment of a control construct without an insulator (noIns) is indicated as a dotted line. In **c**, **e**, **g**, and **j**, the dashed line represents a linear regression line and error bars represent the 95% confidence interval. Pearson’s *R*^2^, Spearman’s *ρ*, and number (*n*) of constructs are indicated.

### Active fragments can be assembled into strong insulators

We asked whether insulator activity can be increased by combining up to three fragments. We selected 26 fragments with high insulator activity (top 25% of all fragments) in tobacco and 6 fragments with low insulator activity (bottom 25% of all fragments) in tobacco (Supplementary Table 1). These fragments were used in the forward and reverse orientation to build constructs with both the individual fragments and with the over 2,900 randomly generated two-fragment combinations. Additionally, we built over 13,000 three-fragment combinations that added one of five fragments with very high insulator activity (top 5% of all fragments; Supplementary Table 1) upstream of the randomly generated two-fragment combinations. Fragments and fragment combinations were cloned between the 35S enhancer and 35S minimal promoter (Fig 3a). Increasing the number of insulator fragments increased insulator activity. In tobacco, most constructs with three insulator fragments completely blocked the 35S enhancer (Fig. 3b and Supplementary Data 2). Combinations of fragments derived from different full-length insulators showed a similar activity distribution to combinations of fragments derived from the same full-length insulator. Similarly, the activity distribution of combinations with two copies of the same fragment was largely indistinguishable from that of combinations with two non-identical fragments.

**Fig. 3.**
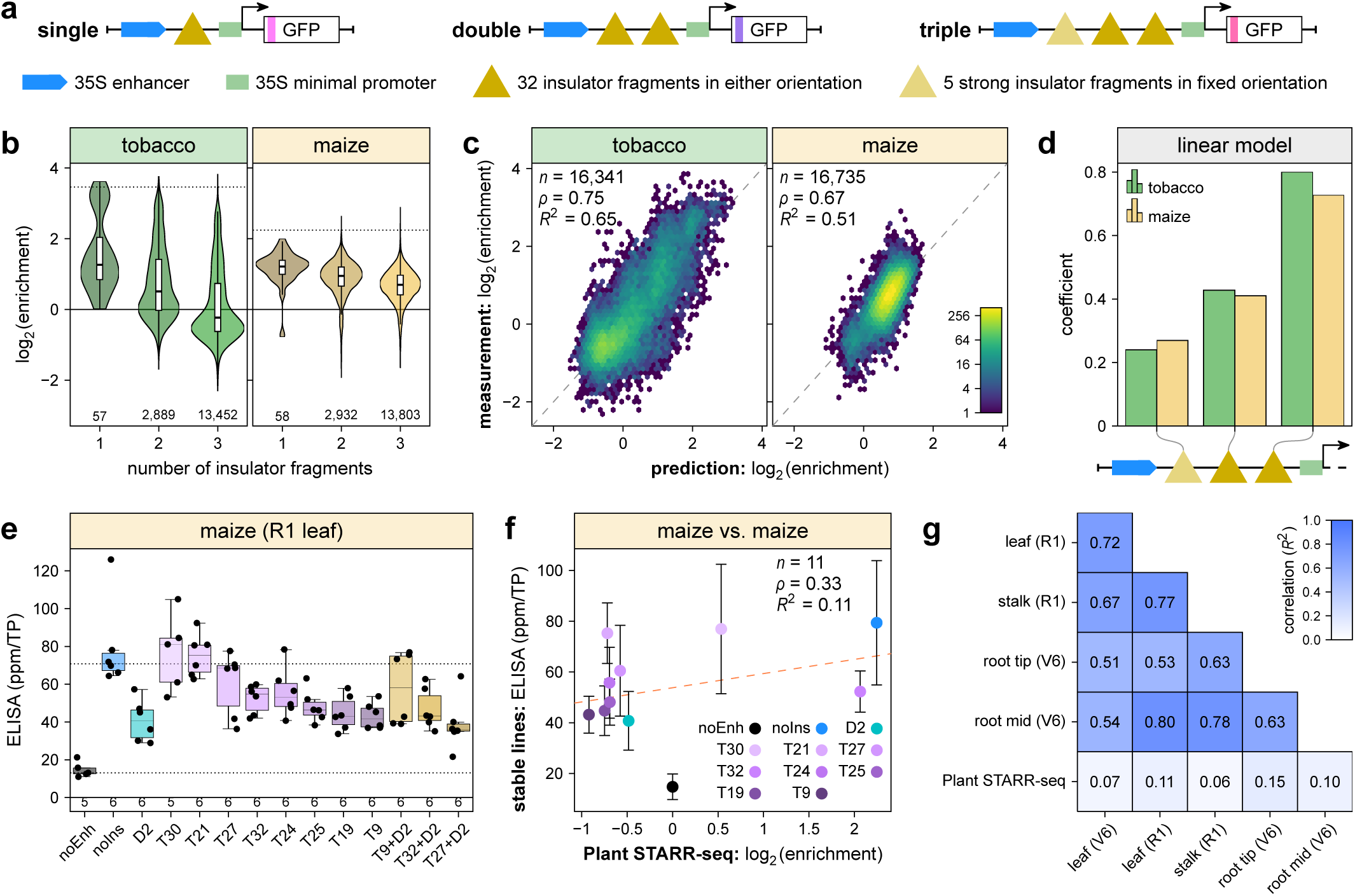
Insulator fragments can be stacked to create very strong enhancer-blocking insulators. **a**, One, two, or three 170-bp fragments of known insulators were cloned between a 35S enhancer and a 35S minimal promoter driving the expression of a barcoded GFP reporter gene. **b**, All insulator constructs were pooled and subjected to Plant STARR-seq in tobacco leaves (tobacco) and maize protoplasts (maize). Reporter mRNA enrichment was normalized to a control construct without an enhancer or insulator (log2 set to 0). Violin plots are as defined in Fig. 1b. The enrichment of a control construct without an insulator is indicated as dotted line. **c**, A linear model was trained to predict the enrichment of stacked insulator constructs based on the activity of individual insulator fragments and their position within the construct. The correlation between the model’s prediction (prediction) and experimentally determined enrichment values (measurement) is shown as a hexbin plot (color represents the count of points in each hexagon; **c**). Pearson’s *R*^2^, Spearman’s *ρ*, and number (*n*) of fragments are indicated. **d**, Coefficients assigned by the linear model to insulator fragments in the indicated positions of the stacked constructs. **e**,**f**, The activity of insulator fragment combinations in constructs as in Fig. 2h was measured in R1 leaves of transgenic maize lines (**e**) and compared to the corresponding results from Plant STARR-seq in maize protoplasts (**f**). **g**, Correlation (Pearson’s *R*^2^) between the expression of all tested constructs across different tissues and developmental stages. The correlation with Plant STARR-seq results from maize protoplasts is also shown.

We trained a linear model based on the insulator activity of the individual fragments and their position in the construct to predict the insulator activity of two-fragment and three-fragment combinations in tobacco and maize (Fig. 3c). Model accuracy was similar for the two-fragment and three-fragment combinations in tobacco (*R*^2^ of 0.67 and 0.62, respectively). In maize, prediction accuracy was higher for the two-fragment combinations than for the three-fragment combinations (*R*^2^ of 0.60 and 0.48, respectively). The model coefficients showed that the fragment closest to the minimal promoter contributes the most to the combined insulator activity, while the fragment closest to the enhancer contributes the least (Fig. 3d). Taken together, the insulator activity of the individual fragments appears to be the key determinant for the activity of the fragment combinations.

Next, we tested the activity of one two-fragment combination and nine three-fragment combinations (Supplementary Table 2) in stable maize plants. Most of these fragment combinations showed insulator activity in the transgenic maize plants (Fig. 3e and Extended Data Fig. 4a). However, their activity was weaker than observed in the Plant STARR-seq experiments, likely because we used a moderate-strength promoter from maize for the transgenic maize reporter constructs instead of the minimal 35S promoter used in Plant STARR-seq (Fig. 3f and Extended Data Fig. 4b). To further increase insulator activity, we cloned the two-fragment combination D2 downstream of the three-fragment combinations T9, T32, and T27 (Supplementary Table 2) to yield three constructs of five fragments (T9+D2, T32+D2, and T27+D2). These five-fragment combinations showed similar insulator activity as the corresponding two- or three-fragment combinations (Fig. 3e and Extended Data Fig. 4a), indicating diminishing returns from stacking increasing numbers of fragments. Because most insulator combinations reached the detection limit in our Plant STARR-seq assay but not in the stable maize plants, the correlation between the ELISA and Plant STARR-seq data was low (Fig. 3f,g and Extended Data Fig. 4b). However, we observed a strong correlation between ELISA results for samples obtained from different plant tissues (leaf, stalk, and root) and developmental stages (V6 and R1). This observation is consistent with our results for single insulator fragments and indicates that insulator activity is not strongly affected by tissue identity or developmental stage.

### Insulator-derived fragments also exhibit silencer activity

The comparison of the Plant STARR-seq and stable maize data suggests that insulator activity might be promoter-dependent. To investigate this hypothesis, we built constructs with hybrid promoters by inserting the *AB80* or *Cab-1* enhancer between the 35S minimal promoter and the insulator fragments and tested if an additional downstream enhancer affected the ability of the insulator-derived fragments to block an upstream 35S enhancer (Fig 4a, top). Many fragments showed insulator activity with both downstream enhancers (Fig. 4b, left and Supplementary Data 3) and this activity was only slightly weaker than in constructs without a downstream enhancer (Extended Data Fig. 5). This finding suggests that the insulator-derived fragments remain active at a greater distance and work with more complex promoters than the short 35S minimal promoter.

**Fig. 4.**
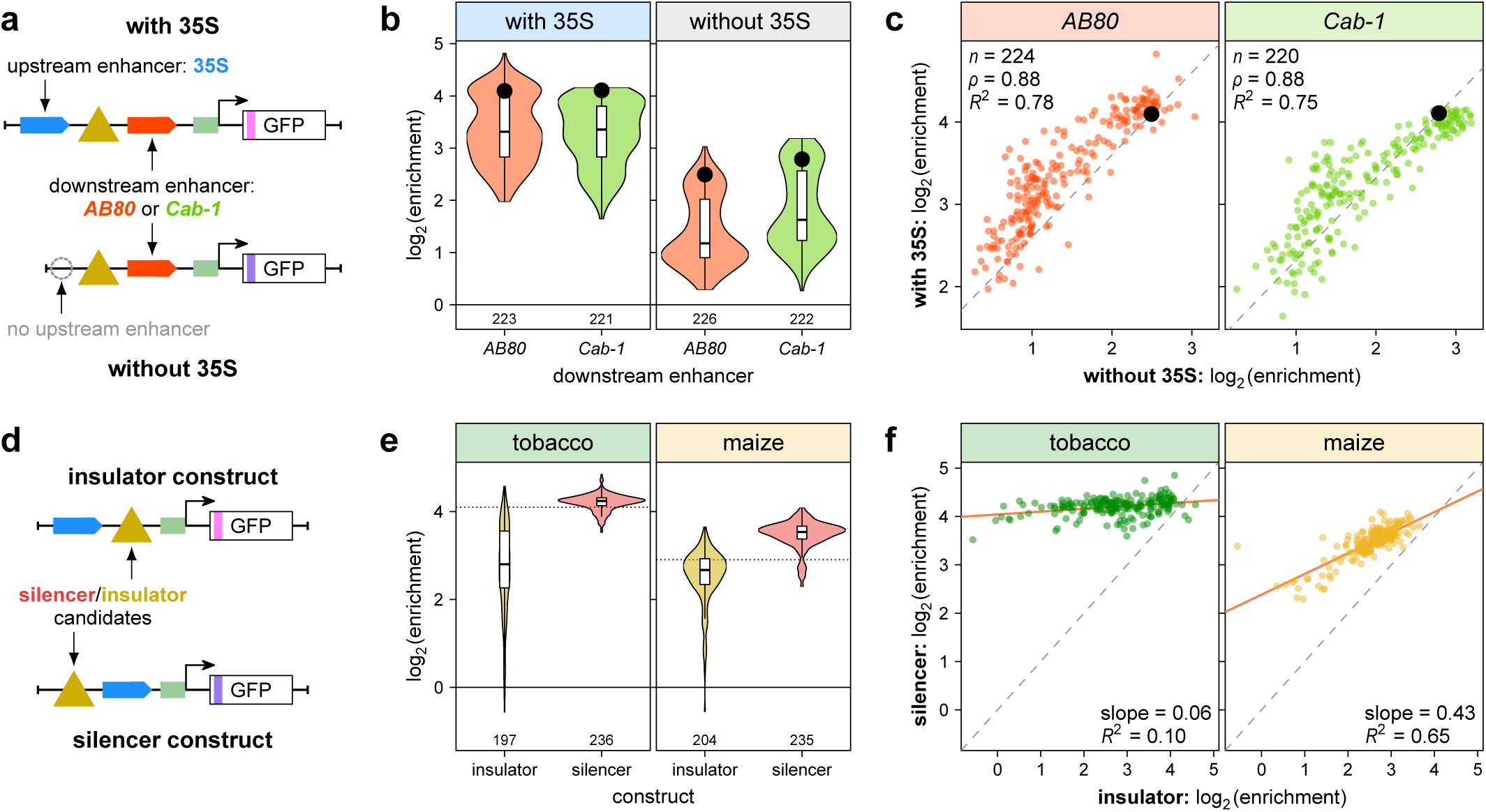
Insulators exhibit silencer activity in some contexts. **a**, Insulator fragments were cloned upstream of a *AB80* or *Cab-1* enhancer and a 35S minimal promoter driving the expression of a barcoded GFP reporter gene. Half of the constructs also harbored a 35S enhancer upstream of the insulator fragments (with 35S) while the other half lacked an upstream enhancer (without 35S). **b**, All constructs were pooled and subjected to Plant STARR-seq in tobacco leaves. Reporter mRNA enrichment was normalized to a control construct without an enhancer or insulator (noEnh; log2 set to 0). The enrichment of a control construct without an insulator is indicated as a black dot. **c**, Correlation between insulator fragment activity in constructs with or without the upstream 35S enhancer. The dashed line represents a y = x line fitted through the point corresponding to a control construct without an insulator (black dot). **d**, Insulator fragments were cloned in between (insulator construct) or upstream of (silencer construct) a 35S enhancer and a 35S minimal promoter driving the expression of a barcoded GFP reporter gene. **e**, All constructs were pooled and subjected to Plant STARR-seq in tobacco leaves (tobacco) or maize protoplasts (maize). Reporter mRNA enrichment was normalized to a control construct without an enhancer or insulator (noEnh; log2 set to 0).The enrichment of a control construct without an insulator is indicated as a black dot. **f**, Comparison of the enrichment of insulator fragments in insulator or silencer constructs. A linear regression line is shown as a solid line and its slope and goodness-of-fit (*R*^2^) is indicated. Violin plots in **b** and **e** are as defined in Fig. 1b.

We also tested a set of control constructs without the upstream 35S enhancer (Fig 4a, bottom) and found that many insulator fragments resulted in lower enrichment than a control construct without an insulator fragment (Fig. 4b, right and Supplementary Data 3), indicating transcriptional repression. The enrichment of reporter constructs with and without the upstream 35S enhancer was well correlated (Fig. 4c). These results demonstrate that fragments derived from characterized insulators and that showed enhancer-blocking activity in Plant-STARR-seq can also function as transcriptional silencers.

To rigorously assess whether the insulator-derived fragments had silencer activity, we built a new library with two different construct layouts: (i) in the ‘insulator’ construct, the fragments were inserted between the 35S enhancer and 35S minimal promoter; and (ii) in the ‘silencer’ construct, the fragments were inserted upstream of the 35S enhancer (Fig. 4d). As before, many fragments led to a reduced enrichment of the reporter gene when inserted between the enhancer and promoter (*i.e.,* in the insulator construct). The insulator-derived fragments showed little to no activity in the silencer construct in tobacco; however, we observed some silencer activity in maize (Fig. 4e and Supplementary Data 4).

We reasoned that the activity of fragments in the insulator construct might be a combination of enhancer-blocking and silencer activity. To quantify what fraction of the apparent insulator activity could be explained by transcriptional repression rather than insulation, we plotted the activities of all fragments in the insulator construct against their activities in the silencer construct (Fig. 4f). The slope of the regression line in these plots is a proxy for the maximal contribution of transcriptional repression to the apparent insulator activity. Up to 6% and 43% of the observed activity in the insulator construct could be explained by silencer activity in tobacco and maize, respectively (Fig. 4f).

### Silencer activity depends on enhancer strength

Because we found evidence of silencer activity in tobacco leaves in constructs containing the *AB80* or *Cab-1* enhancer (Fig. 4b,c), but not in those with the strong 35S enhancer (Fig. 4e,f), we built insulator and silencer constructs with eight different enhancers (Fig. 5a,b). These enhancers showed a wide range of strength in tobacco but were all, apart from the 35S enhancer, weak in maize (Fig. 5b).

**Fig. 5.**
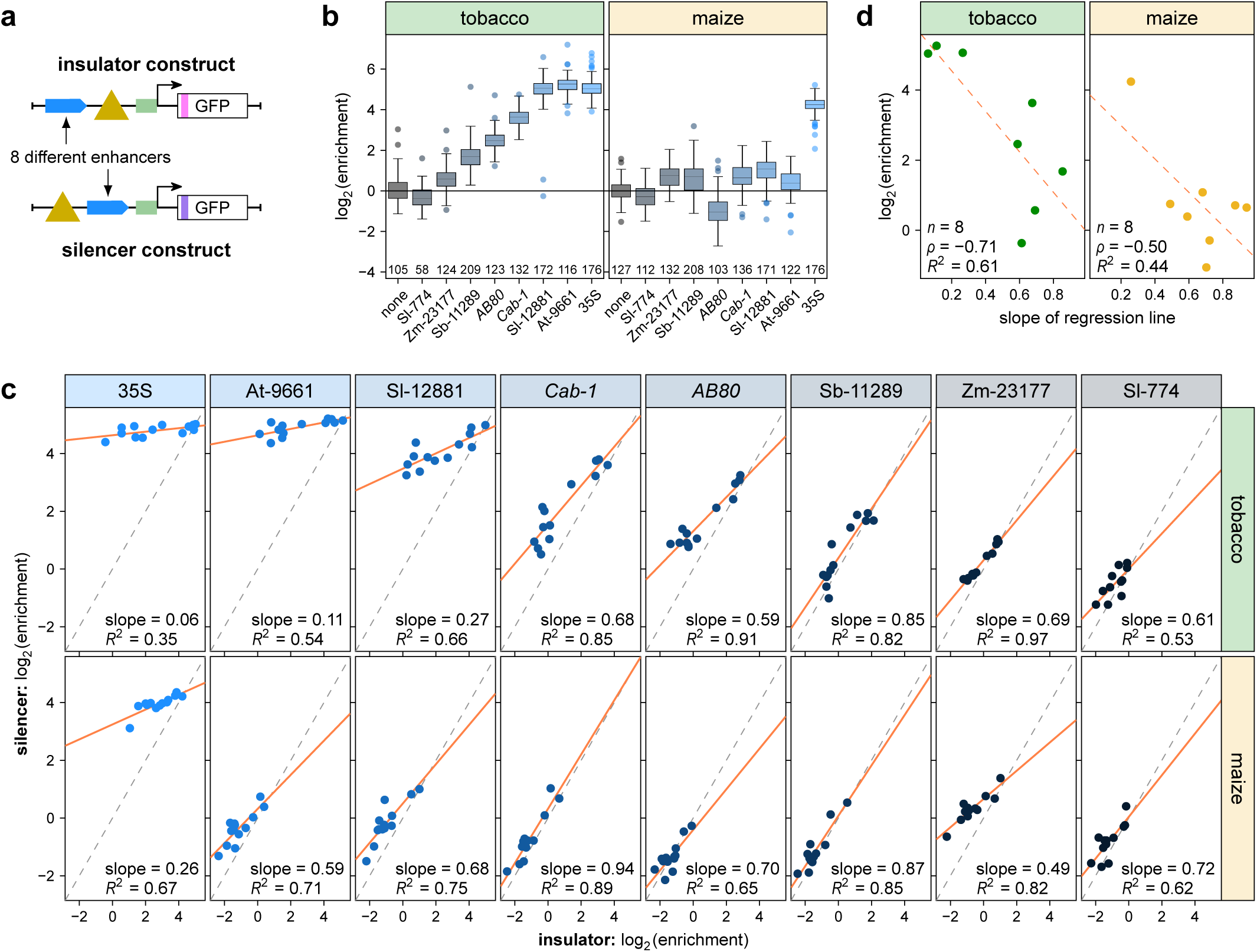
Silencer activity depends on enhancer strength. **a**, Selected insulators and insulator fragments were cloned in between (insulator construct) or upstream of (silencer construct) an enhancer and a 35S minimal promoter driving the expression of a barcoded GFP reporter gene. Eight different enhancers were used to build these constructs. All constructs were pooled and subjected to Plant STARR-seq in tobacco leaves (tobacco) or maize protoplasts (maize). **b**, Strength of the eight enhancers in constructs without an insulator. Reporter mRNA enrichment was normalized to a control construct without an enhancer (none; log2 set to 0). Box plots represent the median (center line), upper and lower quartiles, and 1.5× interquartile range (whiskers) for all corresponding barcodes from two independent replicates. Numbers at the bottom of the plot indicate the number of samples in each group.**c**, Comparison of the enrichment of insulators and insulator fragments in insulator or silencer constructs. A linear regression line is shown as a solid line and its slope and goodness-of-fit (*R*^2^) is indicated. **d**, Correlation between the slope of the regression lines from **c** and the strength of the corresponding enhancer (see **b**). Pearson’s *R*^2^, Spearman’s *ρ*, and number (*n*) of constructs are indicated. A linear regression line is shown as a dashed line.

We tested these enhancers with six full-length insulators and six insulator-derived fragments (Supplementary Table 3). Insulators and insulator fragments showed little activity as silencers with strong enhancers (like the 35S, At-9661, and Sl-12881 enhancers in tobacco and the 35S enhancer in maize) but much more activity as silencers with weak enhancers (Fig. 5c and Supplementary Data 5). This result conclusively demonstrates that these previously identified insulators and their fragments can function as enhancer-blocking insulators or as silencers depending on regulatory context.

As before, we plotted the enrichment of fragments in insulator constructs against their enrichment in silencer constructs. We used the slope of a linear regression line as a proxy to determine how much of the apparent insulator activity could be explained by silencer activity. For constructs with strong enhancers, between 6% and 27% of the apparent insulator activity could be explained by silencer activity. This proportion increased with weak enhancers, such that silencer activity could explain up to 94% of the observed activity in the insulator construct. Overall, the slopes negatively correlated with the strength of the corresponding enhancer (Fig. 5d).

To test whether the insulators showed silencer activity when integrated into the genome, we used dual-luciferase reporter constructs with the insulator residing upstream of the 35S or *AB80* enhancer to generate stable transgenic *Arabidopsis* plants (Fig. 6a). As in the transient Plant STARR-seq experiments, the insulators showed no silencer activity with the strong 35S enhancer and partial silencer activity with the somewhat weaker *AB80* enhancer in transgenic *Arabidopsis* plants (Fig. 6b-d). Taken together, these results are consistent with the observation that previously identified insulators show silencer activity that is inversely correlated with the strength of the enhancer with which they are paired.

**Fig. 6.**
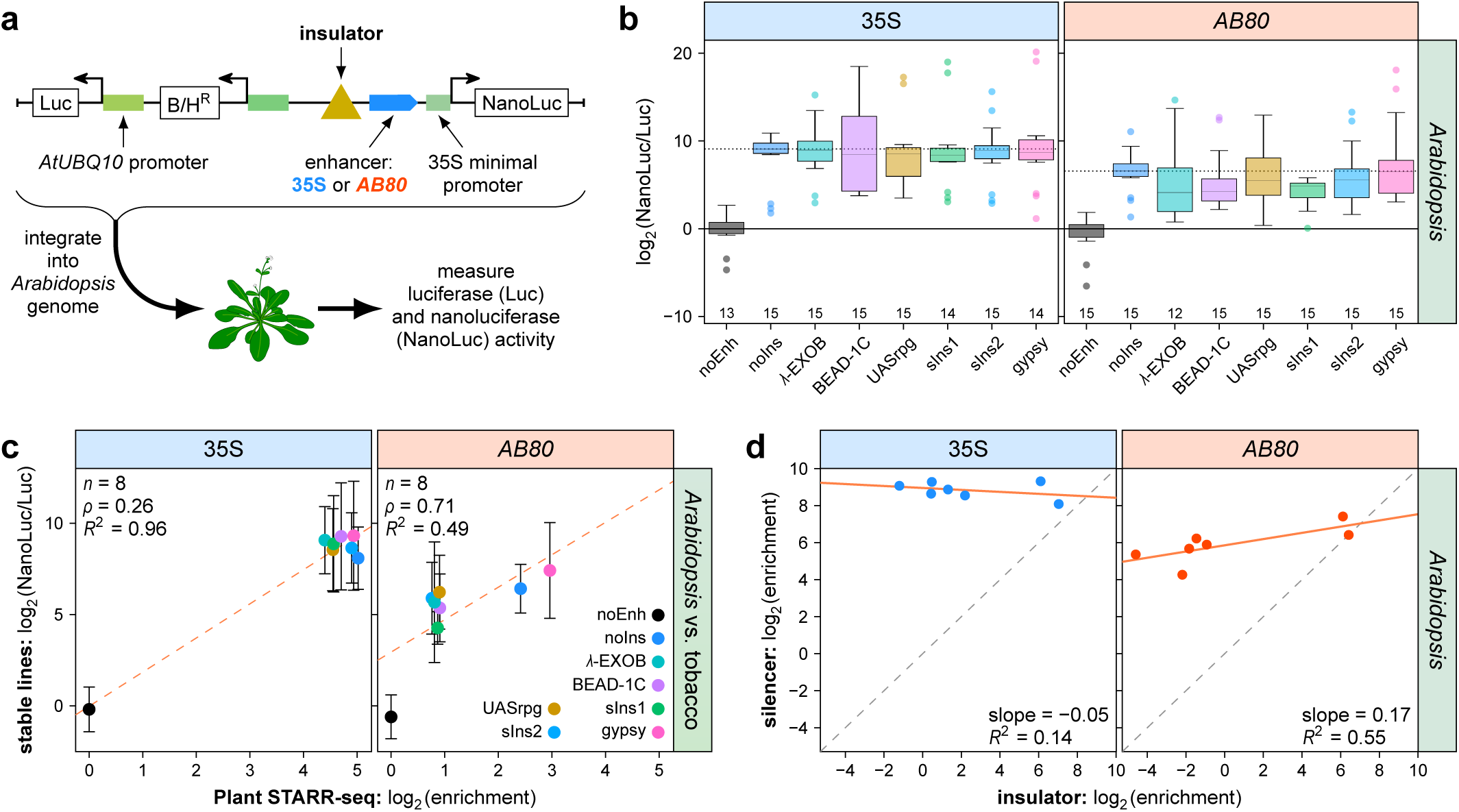
Enhancer-dependent silencer activity in stable transgenic plants. **a**, Transgenic *Arabidopsis* lines were generated with T-DNAs harboring a constitutively expressed luciferase (Luc) gene and a nanoluciferase (NanoLuc) gene under control of a 35S minimal promoter coupled to the 35S or *AB80* enhancer (as indicated above the plots) with insulator candidates inserted upstream of the enhancer. Nanoluciferase activity was measured in at least 4 plants from these lines and normalized to the activity of luciferase. The NanoLuc/Luc ratio was normalized to a control construct without an enhancer or insulator (noEnh; log2 set to 0). **b**,**c**, The activity of full-length insulators was measured in *Arabidopsis* lines (**b**) and compared to the corresponding results from Plant STARR-seq in tobacco leaves (**c**). Box plots in **b** are as defined in Fig. 2. In **c**, the dashed line represents a linear regression line and error bars represent the 95% confidence interval. Pearson’s *R*^2^, Spearman’s *ρ*, and number (*n*) of constructs are indicated. **d**, Comparison of the mean NanoLuc/Luc ratio of full-length insulators in insulator (Fig. 2b) or silencer constructs (**b**). A linear regression line is shown as a solid line and its slope and goodness-of-fit (*R*^2^) is indicated.

## Discussion

Using the high throughput Plant STARR-seq assay on fragments of insulators known to be functional in plants, we identified more than 100 170-bp fragments with enhancer-blocking activity. These short fragments could be combined to generate stronger insulators, some capable of completely blocking the activity of the viral 35S enhancer. The fragments were active as insulators with different enhancers and promoters and across diverse plant tissues. Surprisingly, these insulators and their fragments showed silencer activity when coupled with weak enhancers. Consistent with other work, this finding showcases the complexity of regulatory grammar, wherein *cis*-regulatory elements can have multiple activities that may be observed only in specific conditions or contexts^6^. For example, mesoderm-specific *Drosophila* silencers often function as enhancers in other cell types^22^. Thus, regulatory elements must be tested systematically in different contexts – *e.g.*, as insulators, silencers, or enhancers, and across species and tissues – to understand the mechanistic underpinnings of their potentially complex functions.

The elements studied here behaved like classical enhancer-blocking insulators in combination with strong enhancers, as they reduced reporter expression only when inserted between the enhancer and promoter. In contrast, with weak enhancers, the same elements behaved like typical silencers that repressed transcription in a position-independent manner. With intermediate strength enhancers, a continuum between these two extremes was observed: The elements reduced transcription when placed either upstream or downstream of the enhancer, but the effect was stronger in the downstream context. It remains to be determined if the observed activity is a combination of distinct insulator and silencer functions or if we identified a novel, yet unknown regulatory mechanism. Our observation that, independent of enhancer strength, the activity of fragments in insulator and silencer constructs is well correlated suggests that the latter might be the case.

To date, the molecular mechanisms underlying plant insulator function are unknown. In animals, several DNA-binding proteins, including su(Hw), BEAF-32, and Zw5 in *Drosophila*^27–29^ and CTCF in humans^30^, play a role in insulator function. However, homologs of these proteins have not been identified in plants. The number of fragments with insulator activity tested here is too small to derive putative protein-binding motifs with confidence. Moreover, there is no evidence that insulation in plants requires protein binding. In contrast to enhancer activity^6,31^, we found that insulator activity was orientation-dependent, as has been observed in animals^32,33^ and previously in plants^12,34^. In some cases, orientation-dependence is a consequence of composite elements with both insulator and enhancer activities^12,32,33^. An alternative hypothesis for orientation dependence is that structural properties of the insulator DNA contribute to insulator function. This hypothesis is also consistent with our finding that GC content is a major contributor to insulator activity.

Short insulators elements are useful for plant biotechnology to minimize the size of transgene cassettes to ensure efficient transformation. Transgene cassettes, especially those composed of multiple genes, often show unpredictable expression patterns even when the regulation of the individual genes is well-characterized. The insulators identified here are promising building blocks to make expression more predictable and thus plant engineering more economically feasible. Insulator activity showed some specificity to the tobacco or maize system, suggesting that insulators need to be designed for either dicots or monocots. Although our work shows that the use of insulators in transgene cassettes must account for both their silencer and insulator activities, plant biotechnology efforts tend to use strong constitutive promoters, such that silencer activity is negligible. Moreover, when used with tissue- or condition-specific enhancers, insulators with enhancer-dependent silencer activity could be beneficial.

Such insulator-enhancer combinations could repress leaky expression in tissues or conditions in which the enhancer is inactive and insulate expression when the enhancer becomes fully active. Similarly, the dual-function elements identified here might be used to fine-tune transgene expression by repressing overly-active transcription while simultaneously isolating the transgene from other surrounding regulatory elements.

## Methods

### Library design and construction

The full-length λ-EXOB, BEAD-1C, UASrpg, and gypsy insulators were ordered as synthesized DNA fragments. The synthetic insulators sIns1 and sIns2 were PCR amplified from pZS*11_4enh (Addgene no. 149423; https://www.addgene.org/149423/; ref. ^3^) and pEvolvR-enCas9-PolI3M-TBD (Addgene no. 113077; https://www.addgene.org/113077/; ref. ^35^), respectively. Insulator fragments were ordered as an oligonucleotide array from Twist Bioscience with 15-bp flanking sequences for amplification. The 35S, *AB80*, and *Cab-1* enhancers were PCR amplified from pZS*11_4enh. The At-9661, Sl-12881, Sb-11289, Zm-23177, and Sl-774 enhancers were ordered as synthesized DNA fragments. The sequences of the full-length insulators and the oligonucleotides used in this study are listed in Supplementary Table 4 and Supplementary Table 5, respectively.

All libraries used in this study were constructed using pPSm, a shortened version of pPSup (Addgene no. 149416; https://www.addgene.org/149416/; ref. ^3^) lacking the BlpR cassette, as the base plasmid. The plasmid’s T-DNA region harbors a GFP reporter construct terminated by the poly(A) site of the Arabidopsis (*Arabidopsis thaliana*) ribulose bisphosphate carboxylase small chain 1A gene. Two versions of pPSm were created to receive insulators in the forward (pPSmF) or reverse (pPSmR) orientation by changing the BsaI scars to ACTC and CTGT or ACAG and GAGT, respectively. The plasmids were deposited at Addgene (Addgene no. 226912 and 226913; https://www.addgene.org/226912/; https://www.addgene.org/226913/). Gibson assembly^36^ was used to insert enhancers into pPSm plasmids. The 35S minimal promoter followed by the 5′ UTR from a maize histone H3 gene (Zm00001d041672), an ATG start codon and a 18-bp random barcode (VNNVNNVNNVNNVNNVNN; V = A, C, or G) was cloned in front of the second codon of GFP by Golden Gate cloning^37^ using BbsI-HF (NEB). To distinguish between sub-libraries, positions 1, 4, 7, 10, 13, and 16 of the barcodes were set to fixed bases. Insulators and insulator fragments were inserted into the pPSm plasmids by Golden Gate cloning using BsaI-HFv2 (NEB). The resulting libraries were bottlenecked to yield about 20–50 barcodes per enhancer.

The base plasmid for dual-luciferase constructs was derived from pDL (Addgene no. 208978; https://www.addgene.org/208978/; ref. ^23^) by changing the BsaI scars to ACTC and CTGT. The 35S or *AB80* enhancer was inserted into this plasmid upstream or downstream of the BsaI Golden Gate cassette via Gibson assembly. Full-length insulators and insulator fragments were inserted by Golden Gate cloning using BsaI-HFv2 (NEB). For rice dual-luciferase constructs, the BlpR cassette was replaced by a hygromycin resistance gene under control of the switchgrass polyubiquitin 2 promoter and the 35S terminator derived from plasmid JD633 (Addgene no. 160393; https://www.addgene.org/160393/; ref. ^38^).

The expression cassettes for the Agrobacterium-based transformation vectors to generate transgenic corn plants consisted of a reporter gene driven by a moderate-strength constitutive promoter coupled to a heterologous intron with either the CaMV 35S enhancer upstream of the promoter (negative control) or no enhancer (positive control). The same terminator was used in cassettes to terminate transcription. The insulators were tested using the expression cassette with the 35S enhancer by inserting them between the 35S enhancer and the promoter.

### Tobacco cultivation and transformation

Tobacco (*Nicotiana benthamiana*) was grown in soil (Sunshine Mix no. 4) at 25°C in a long-day photoperiod (16 h light and 8 h dark; cool-white fluorescent lights [Philips TL-D 58 W/840]; intensity 300 μmol m^−2^ s^−1^). Plants were transformed approximately 3 weeks after germination. For transient transformation of tobacco leaves, Plant STARR-seq libraries were introduced into *Agrobacterium tumefaciens* strain GV3101 (harboring the virulence plasmid pMP90 and the helper plasmid pMisoG) by electroporation. An overnight culture of the transformed *A. tumefaciens* was diluted into 100 ml YEP medium (1% [w/v] yeast extract and 2% [w/v] peptone) and grown at 28°C for 8 h. A 5-ml input sample of the cells was collected, and plasmids were isolated from it using the QIAprep Spin Miniprep Kit (QIAGEN) according to the manufacturer’s instructions. The remaining cells were harvested and resuspended in 100 ml induction medium (M9 medium [3 g/L KH_2_PO_4_, 0.5 g/L NaCl, 6.8 g/L Na_2_HPO_4_, and 1 g/L NH_4_Cl] supplemented with 1% [w/v] glucose, 10 mM MES, pH 5.2, 100 μM CaCl_2_, 2 mM MgSO_4_, and 100 μM acetosyringone). After overnight growth, the *Agrobacteria* were harvested, resuspended in infiltration solution (10 mM MES, pH 5.2, 10 mM MgCl_2_, 150 μM acetosyringone, and 5 μM lipoic acid) to an optical density (OD) of 1 and infiltrated into leaves 3 and 4 of two (full-length insulator library) or four (all other libraries) tobacco plants. The plants were further grown for 48 h under normal conditions (16 h light and 8 h dark) or in the dark before mRNA extraction.

### Maize cultivation and transformation

For Plant STARR-seq in maize (*Zea mays* L. cultivar B73), we used PEG transformation method as previously described^39^. Maize seeds were germinated in soil at 25°C in a long-day photoperiod (16 h light and 8 h dark; cool-white fluorescent lights [Philips TL-D 58 W/840]; intensity 300 μmol m^−2^ s^−1^). After 3 days, the seedlings were moved to complete darkness at 25°C and grown for 10–11 days. From each seedling, 10 cm sections from the second and third leaf were cut into thin 0.5 mm strips perpendicular to veins and immediately submerged in 10 ml of protoplasting enzyme solution (0.6 M mannitol, 10 mM MES pH 5.7, 15 mg/ml cellulase R10, 3 mg/ml macerozyme, 1 mM CaCl_2_, 0.1% [w/v] BSA, and 5 mM beta-mercaptoethanol). The mixture was covered in foil to keep out light, vacuum infiltrated for 3 min, and incubated on a shaker at 40 rpm for 2.5 hours. Protoplasts were released by incubating an extra 10 min at 80 rpm. To quench the reaction, 10 mL ice-cold MMG (0.6 M Mannitol, 4 mM MES pH 5.7, 15 mM MgCl_2_) was added to the enzyme solution and the whole solution was filtered through a 40 µM cell strainer. To pellet protoplasts, the filtrate was split into equal volumes of no more than 10 mL in chilled round-bottom glass centrifuge vials and centrifuged at 100 x g for 4 min at room temperature (RT). Pellets were resuspended in 1 mL cold MMG each and combined into a single round-bottom vial. To wash, MMG was added to make a total volume of 5 mL and the solution was centrifuged at 100 x g for 3 min at RT. This wash step was repeated two more times. The final pellet was resuspended in 1–2 mL of MMG. A sample of the resuspended protoplasts was diluted 1:20 in MMG and used to count the number of viable cells using Fluorescein Diacetate as a dye. For each replicate, one to ten million protoplasts were mixed with 15–150 µg of the Plant STARR-seq plasmid library in a fresh tube, topped with MMG to a volume of 114.4 µL per million protoplasts, and incubated on ice for 30 min. For PEG transformation, 105.6 µL per million protoplasts of PEG solution (0.6 M Mannitol, 0.1 M CaCl_2_, 25% [w/v] poly-ethylene glycol MW 4000) was added to reach a final concentration of 12% (w/v) PEG. The mixture was incubated for 10 min in the dark at RT. After incubation, the transformation solution was diluted with five volumes incubation solution (0.6 M Mannitol, 4 mM MES pH 5.7, 4 mM KCl), and centrifuged at 100 x g for 4 min at RT. The protoplast pellet was washed with 5 mL of incubation solution, centrifuged at 100 x g for 3 min at RT, and resuspended in incubation solution to a concentration of 500 cells/µL. Protoplasts were incubated overnight in the dark at RT to allow for transcription of the plasmid library and then pelleted (4 min, 100 x g, RT). The pellet was washed with 1–5 mL incubation solution and centrifuged (3 min, 100 x g, RT). The pellet was finally resuspended in 1–5 mL incubation solution. An aliquot of the solution was used to check transformation efficiency under a microscope. Cells were pelleted (4 min, 100 x g, RT) and resuspended in 1–2 mL Trizol for subsequent mRNA extraction. An aliquot of the plasmid library used for PEG transformation was used as the input sample for Plant STARR-seq.

To generate stable transgenic maize plants, we followed a previously published procedure^40^.

### Arabidopsis cultivation and transformation

*Arabidopsis thaliana* Col-0 was grown in soil (Sunshine Mix no. 4) at 20°C in a long-day photoperiod (16 h light and 8 h dark; cool-white fluorescent lights [Sylvania FO32/841/ECO 32W]; intensity 100 μmol m^−2^ s^−1^). For transformation, dual-luciferase plasmids were introduced into *Agrobacterium tumefaciens* strain GV3101 (harboring the virulence plasmid pMP90 and the helper plasmid pMisoG) by electroporation. Transgenic *Arabidopsis* plants were generated by floral dipping^41^ and selected for by spraying with a 0.01% (w/v) Glufosinate solution.

### Rice cultivation and transformation

The rice (*Oryza sativa* L. ssp. *japonica*) cultivar Kitaake was used for genetic transformation following a previously described protocol^42^ with slight modifications. The mature seeds were sterilized with a 7.5% (w/v) sodium hypochlorite solution for 20 minutes, followed by three sterile water rinses. The seeds were placed on callus induction medium (4.4 g/L MS salts with vitamins, 30 g/L sucrose, 2 mg/L 2,4-dichlorophenoxyacetic acid, 8 g/L agar, pH 5.8) to induce callus cells from scutellum for 10 days. The calli were co-cultivated on callus induction medium supplemented with 200 µM of acetosyringone for 3 days with the *Agrobacterium* strain EHA101 (OD = 0.5) carrying individual insulator constructs. The callus cells were transferred to callus induction medium supplemented with 300 mg/L timentin and 50 mg/L hygromycin for two rounds of selection. The hygromycin resistant callus cells of individual lines were transferred to regeneration medium (4.4 g/L MS salts with vitamins, 30 g/L sucrose, 3 mg/L 6-benzylaminopurine, 0.5 mg/L 1-naphthaleneacetic acid, 8 g/L agar, 25 mg/L hygromycin, 150 mg/L timentin, pH 5.8) for about two rounds to regenerate shoots. The shoots were transferred to rooting medium (4.4 g/L MS salts with vitamins, 30 g/L sucrose, 25 mg/L hygromycin, 8 g/L agar, pH 5.8) and were grown till healthy roots were produced before transferring to soil. The plantlets were transferred to a plastic box containing topsoil from the research farm at the University of Missouri flooded with water. The plantlets were grown in a greenhouse with a short-day photoperiod (12 h light and 12 h dark) at 28°C and 24°C during the day and night, respectively.

### Plant STARR-seq

For all tobacco Plant STARR-seq experiments, two independent biological replicates were performed. Different plants and fresh *Agrobacterium* cultures were used for each biological replicate.

Tobacco leaves were harvested 2 days after infiltration and partitioned into batches of 4 leaves. The leaf batches were frozen in liquid nitrogen, finely ground with mortar and pestle, and immediately resuspended in 10 mL QIAzol (Qiagen). The suspensions were cleared by centrifugation (5 min, 4,000 x g, 4°C). The supernatant was transferred to a 15 mL MaXtract High Density tube (Qiagen) and mixed with 2.5 mL chloroform. After centrifugation (10 min, 1,000 x g, 4°C), the supernatant (approximately 7 mL) was poured into a new tube, and mixed by inversion with 3.5 mL high salt buffer (0.8 M sodium citrate, 1.2 M NaCl) and 3.5 mL isopropanol. The solution was incubated for 15 min at RT to precipitate the RNA and centrifuged (30 min, 4,000 x g, 4°C). The pellet was washed with 10 mL ice-cold 70% ethanol, centrifuged (5 min, 4000 x g, 4°C), and air-dried. The pellet was resuspended in 625 µL of warm (65°C) nuclease-free water and transferred to a new tube. The solution was supplemented with 70 µL 20X DNase I buffer (1 mM CaCl_2_, 100 mM Tris pH 7.4), 70 µL 200 mM MnCl_2_, 5 µL DNase I (ThermoFisher Scientific), and 1 µL RNaseOUT (ThermoFisher Scientific). After 1 h incubation at 37°C, the reaction was stopped with 50 µL 500 mM EDTA. To precipitate the RNA, 375 µL high salt buffer and 375 µL isopropanol were added. After incubation for 15 min at room RT, the RNA was pelleted by centrifugation (20 min, 20,000 x g, 4°C). The pellet was washed with 1 mL ice-cold 70% ethanol, centrifuged (5 min, 20,000 x g, 4°C), air-dried, and resuspended in 50 µL nuclease-free water. All batches of the same sample were pooled, and the solution was supplemented with 0.5 µL RNaseOUT. For cDNA synthesis, two to four reactions with 11 µL RNA solution, 1 µL 10 µM GFP-specific reverse transcription primer, and 1 µL 10 mM dNTPs were incubated at 65°C for 5 min then immediately placed on ice. The reactions were supplemented with 4 µL 5X SuperScript IV buffer, 1 µL 100 mM DTT, 1 µL RNaseOUT, and 1 µL SuperScript IV reverse transcriptase (ThermoFisher Scientific). To ensure that the samples were largely free of DNA contamination, four reactions were used as controls, where the reverse transcriptase and RNaseOUT were replaced with water. Reactions were incubated for 10 min at 55°C, followed by 10 min at 80°C. Sets of 4 reactions each were pooled. The cNDA was purified with the Clean&Concentrate-5 kit (Zymo Research), and eluted in 20 µL 10 mM Tris. The barcode was amplified with 10-20 cycles of polymerase chain reaction (PCR) and read out by next generation sequencing.

For Plant STARR-seq in maize protoplasts, the protoplast-containing Trizol solution from PEG transformation was transferred to 2 mL Phasemaker tubes (1 mL per tube; ThermoFisher Scientific), mixed thoroughly with 300 µL chloroform, and centrifuged (5 min, 15,000 x g, 4°C). RNA was extracted using the RNeasy Plant Mini Kit (QIAGEN). The supernatant was transferred to a QIAshredder column and centrifuged (2 min, 20,000 x g, RT). The flowthrough was transferred to a new 1.5 mL tube and mixed with 300 µL 100% ethanol. Up to 500 µL of the solution was loaded on an RNeasy mini spin column. After centrifugation (10 seconds, 16,100 x g, RT) the flowthrough was discarded. This was repeated until the whole solution had been added to the column. The column was washed with 350 µL RW1 buffer followed by centrifugation (30 sec, 16,100 x g, RT). An on-column DNase I digestion was performed with 70 µL RDD buffer and 10 µL DNase I (Qiagen) for 15 min at RT. The column was washed once with 350 µL RW1 buffer and twice with 500 µL RPE buffer. After each wash step, the column was centrifuged (30 sec, 16,100 x g, RT) and the flowthrough was discarded. The column was dried with an extra centrifugation step (30 sec, 16,100 x g, RT) and transferred to a 1.5 mL collection tube. For elution, 50 µL of RNase-free water was added, and the column was incubated for 1 min, and centrifuged (1 min, 16,100 x g, RT). This elution step was repeated with an additional 40 µL of RNase-free water. The eluate was treated with DNase I (5 µL of 20x DNaseI buffer, 5 µL 200 mM MnCl_2_, 1 µL RNaseOUT, and 2 µL DNase I) for 1 h at 37°C. The solution was supplemented with 20 µL 500 mM EDTA, 1 µL 20 mg/mL glycogen, 12 µL ice-cold 8M LiCl, and 300 µL ice-cold 100% ethanol. The solution was incubated 15 min at −80°C, centrifuged (20 min, 20,000 x g, 4°C). The pellet was washed with 500 µL ice-cold 70% ethanol, and centrifuged (3 min, 20,000 x g, 4°C). The pellet was air-dried and resuspended in 100 µL RNase-free water. Reverse transcription, purification, PCR amplification and sequencing were performed as for the tobacco samples.

### Subassembly and barcode sequencing

Paired-end sequencing on an Illumina NextSeq 550 or 2000 platform was used to link insulator fragments to their respective barcodes. The insulator region was sequenced using paired reads (100–150 bp), and two 18-bp indexing reads were used to sequence the barcodes. The paired insulator fragment and barcode reads were assembled using PANDAseq (version 2.11; ref. ^43^). Insulator fragment-barcode pairs with less than 5 reads and insulator fragments with a mutation or truncation were discarded.

For each Plant STARR-seq experiment, barcodes were sequenced using paired-end reads on an Illumina NextSeq 550 or 2000 system. The paired barcode reads were assembled using PANDAseq.

### Computational methods

For analysis of the Plant STARR-seq experiments, the reads for each barcode were counted in the input and cDNA samples. Barcode counts below 5 were discarded. Barcode counts were normalized to the sum of all counts in the respective sample. For barcodes, enrichment was calculated by dividing the normalized barcode counts in the cDNA sample by that in the corresponding input sample. The sum of the normalized counts for all barcodes associated with a given insulator or insulator fragment were used to calculate its enrichment. For each replicate, the enrichment was normalized to the median enrichment. The mean enrichment across all replicates was normalized to the control construct with enhancer or insulator (noEnh) and used for all analyses. Spearman and Pearson’s correlation were calculated using base R (version 4.3.1). Linear regression analysis was performed using the lm() function in base R.

To predict the enrichment of insulator fragment combinations, a liner model was fitted to Plant STARR-seq data using the lm() function in R with the formula: log2(insulator activity) = log2(insulator activity fragment 3) + log2(insulator activity fragment 2) + log2(insulator activity fragment 1), where log2(insulator activity fragment 1–3) is the enhancer strength of the corresponding fragment when tested individually. Fragments are numbered by increasing distance from the minimal promoter (fragment 1 is the fragment closest to the promoter, fragment 3 the most distal one). Insulator activity was calculated with: log2(insulator activity) = log2(enrichment noIns control) - log2(enrichment insulator). For constructs with one or two fragments, log2(insulator activity) was set to 0 for fragments 3 (two-fragment constructs) or 2 and 3 (one-fragment constructs).

### Dual-luciferase assay

Transgenic *Arabidopsis* lines (T2 generation) with dual-luciferase constructs were grown in soil for 3 weeks. A cork borer (4 mm diameter) was used to collect a total of 4 leaf discs from the third and fourth leaf of the plants. The leaf discs were transferred to 1.5 mL tubes filled with approximately 10 glass beads (1 mm diameter), snap-frozen in liquid nitrogen, and disrupted by shaking twice for 5 sec in a Silamat S6 (Ivoclar) homogenizer. The leaf disc debris was resuspended in 100 µL 1X Passive Lysis Buffer (Promega). The solution was cleared by centrifugation (5 min, 20,000 x g, RT) and 10 µL of the supernatant were mixed with 90 µL 1X passive lysis buffer. Luciferase and nanoluciferase activity were measured on a Biotek Synergy H1 plate reader using the Promega Nano-Glo Dual-Luciferase Reporter Assay System according to the manufacturer’s instructions. Specifically, 10 µL of the leaf extracts were combined with 75 µL ONE-Glo EX Reagent, mixed for 3 min at 425 rpm, and incubated for 2 min before measuring luciferase activity. Subsequently, 75 µL NanoDLR Stop&Glo Reagent were added to the sample. After 3 min mixing at 425 rpm and 12 min incubation, nanoluciferase activity was measured. Two independent biological replicates were performed.

For transgenic rice lines with dual-luciferase constructs, 10–15 mg leaf tissue from 3-week old T0 plants was collected in 1.5 mL tubes filled with approximately 10 glass beads (1 mm diameter). The material was snap-frozen in liquid nitrogen and disrupted by shaking twice for 5 sec in a Silamat S6 (Ivoclar) homogenizer. The leaf debris was resuspended in 200 µL 1X Passive Lysis Buffer (Promega). The solution was cleared by centrifugation (5 min, 20,000 x g, RT) and 10 µL of the supernatant were mixed with 90 µL 1X passive lysis buffer. Luciferase and nanoluciferase activity were measured on a Biotek Synergy H1 in the same way as for *Arabidopsis* samples. Two independent technical replicates (using new samples from the same plants as in the first replicate) were performed.

### ELISA

Insulator activity was detected using a quantitative enzyme linked immunosorbent assay (ELISA) on leaf, stalk, silk, and husk tissues collected from transgenic corn plants. Tissue samples were extracted with 0.60-2.5 ml of buffer comprised of phosphate buffered saline containing polysorbate 20 (8.10 mM PBS + 0.05% polysorbate). Extracted samples were centrifuged and the supernatants used for analysis. 96-well plates pre-coated with reporter-specific monoclonal antibody were incubated with standards and the samples (1hr). After incubation and washing, a second reporter specific monoclonal antibody, conjugated to a horseradish peroxidase enzyme (HRP) was added to the plate and incubated (1hr). After incubation, the plates were washed 5 times and the bound protein-antibody complex was detected by adding TMB (3,3’,5,5’-tetramethylbenzidine) substrate which generated a colored product in the presence of HRP. The reaction was stopped by adding an acid solution and the optical density of each well was determined using a plate reader at 450nm. For each plate a standard curve was included. Adjusted sample concentration values were converted from ng mL-1 to ng mg-1 total extractable protein.

## Data availability

The raw sequencing data underlying this article are available in the National Center for Biotechnology Information (NCBI) Sequence Read Archive at http://www.ncbi.nlm.nih.gov/bioproject/1160710. The processed data underlying this article are available on GitHub at https://github.com/tobjores/Small-DNA-elements-that-act-as-both-insulators-and-silencers-in-plants.

## Code availability

The code used for the analysis and to generate the figures is available on GitHub at https://github.com/tobjores/Small-DNA-elements-that-act-as-both-insulators-and-silencers-in-plants.

## Acknowledgements

This work was supported by the National Science Foundation (RESEARCH-PGR grant no. 1748843 to S.F. and C.Q. and PlantSynBio grant no. 2240888 to C.Q.), the German Research Foundation (DFG; postdoctoral fellowship no. 441540116 to T.J., Emmy Noether program grant no. 517938232 to T.J., and Germany’s Excellence Strategy - EXC-2048/1 - project ID 390686111 to T.J.), the National Institutes of Health (T32 training grant no. HG000035 to J.T., NIGMS grant no. R01-GM079712 to J.T.C. and C.Q., and NIGMS MIRA grant no. 1R35GM139532 to C.Q.), and the United States Department of Agriculture (NIFA postdoctoral fellowship no. 2023-67012-39445 to N.A.M.). We also acknowledge Scott Betts, Hyeon-Je Cho, Megan Christenson, Terry Hu, Albert Lu, Leanne Thompson, Kelli Van Waus, Ning Wang, Emily Wu for their contributions to the work.

## Author contributions

All authors conceived and interpreted experiments; T.J., N.A.M., J.T., S.N.C., BB.L., V.G.-A., and S.J. performed experiments; T.J. analyzed the data and prepared figures; T.J., N.A.M., S.F., and C.Q. wrote the manuscript. All authors read and revised the manuscript.

---

**Extended Data Fig. 1.**
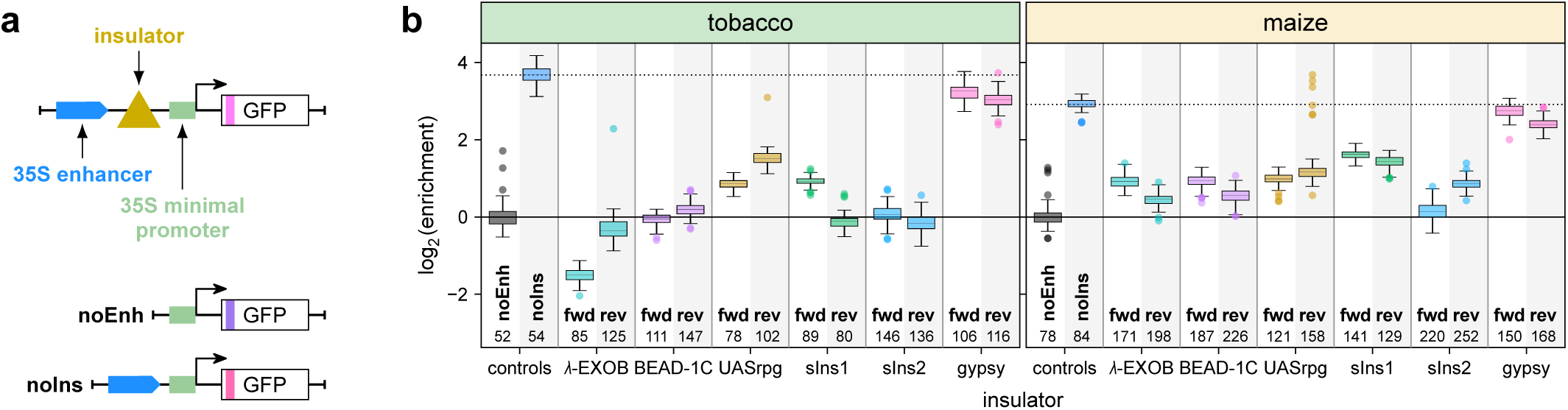
Plant STARR-seq detects activity of enhancer-blocking insulators. **a**, Full-length insulators were cloned in the forward (fwd) or reverse (rev) orientation between a 35S enhancer and a 35S minimal promoter driving the expression of a barcoded GFP reporter gene. **b**, All insulator constructs were pooled and subjected to Plant STARR-seq in tobacco leaves (tobacco) and maize protoplasts (maize). Reporter mRNA enrichment was normalized to a control construct without an enhancer or insulator (noEnh; log2 set to 0). Box plots represent the median (center line), upper and lower quartiles, and 1.5× interquartile range (whiskers) for all corresponding barcodes from two independent replicates. Numbers at the bottom of the plot indicate the number of samples in each group. The enrichment of a control construct without an insulator (noIns) is indicated as a dotted line.

**Extended Data Fig. 2.**
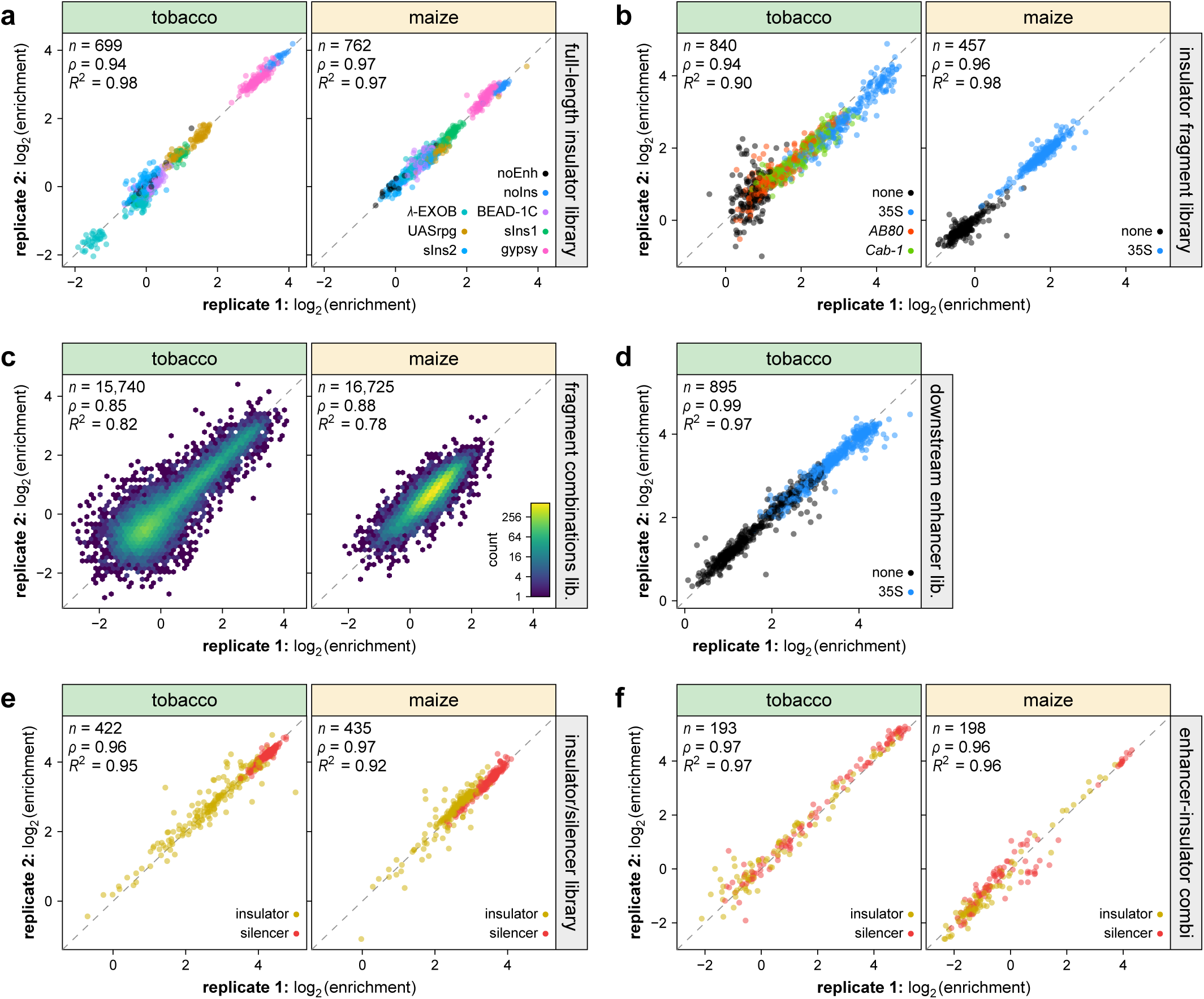
Plant STARR-seq yields highly reproducible results. **a**–**g**, Correlation between biological replicates of Plant STARR-seq for the full-length insulator library used in Extended Data Fig. 1 (**a**), the insulator fragment library used in Fig. 1 (**b**), the insulator fragment combination library used in Fig. 3 (**c**), the downstream enhancer library (**d**) and the insulator/silencer library (**e**) used in Fig. 4, and the enhancer-insulator combination library used in Fig. 5 (**f**). Experiments were performed in tobacco leaves (tobacco) or maize protoplasts (maize) as indicated. Pearson’s *R*^2^, Spearman’s *ρ*, and number (*n*) of constructs are indicated. The color in the hexbin plots in **c** represents the count of points in each hexagon.

**Extended Data Fig. 3.**
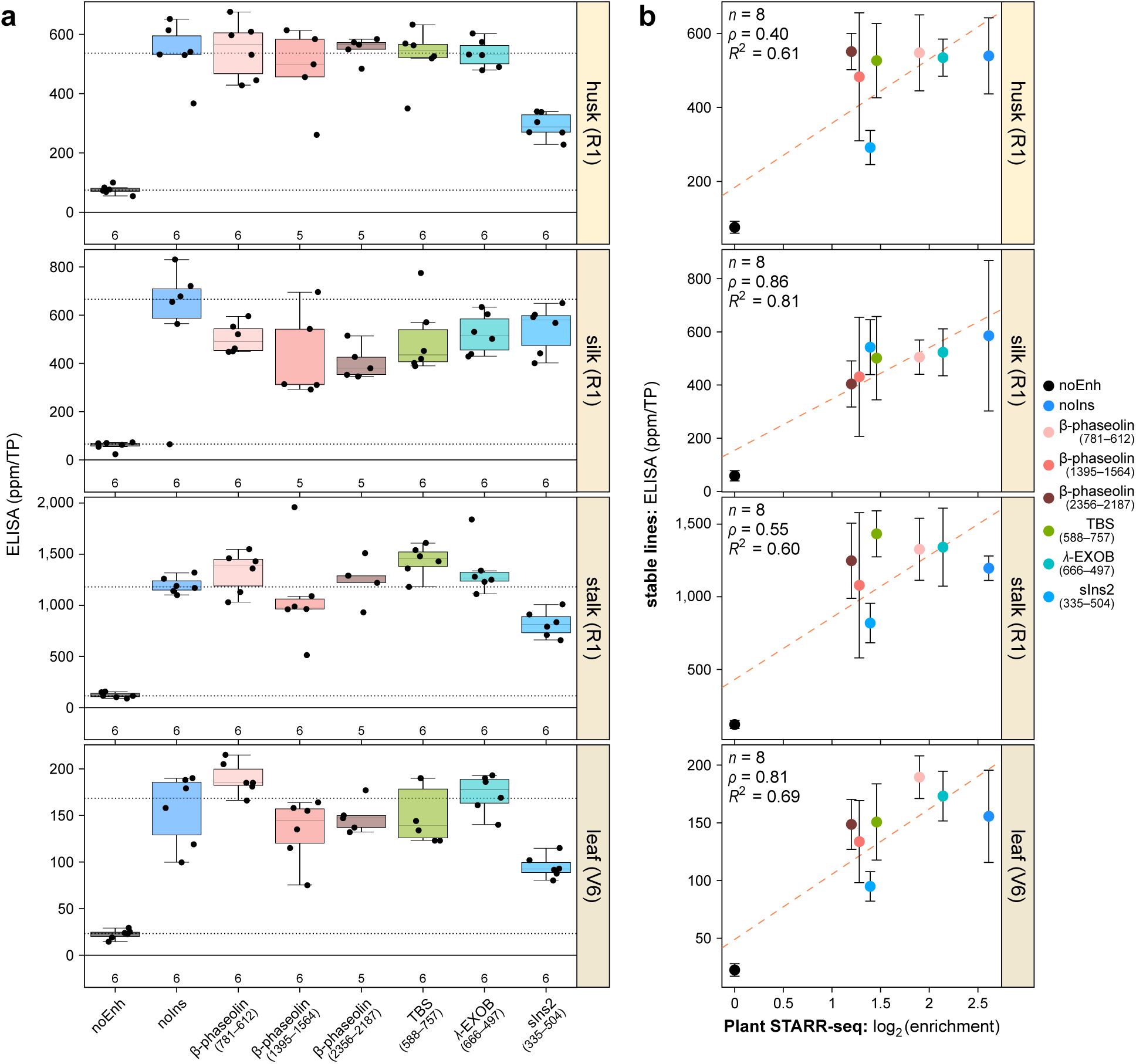
Activity of insulator fragments in different maize tissues. **a**,**b**, Transgenic maize lines were created using constructs as in Fig. 2h. The activity of insulator fragments was measured in the indicated tissues (**a**) and compared to the corresponding results from Plant STARR-seq in maize protoplasts (**b**). Box plots in **a** are as defined in Fig. 2.

**Extended Data Fig. 4.**
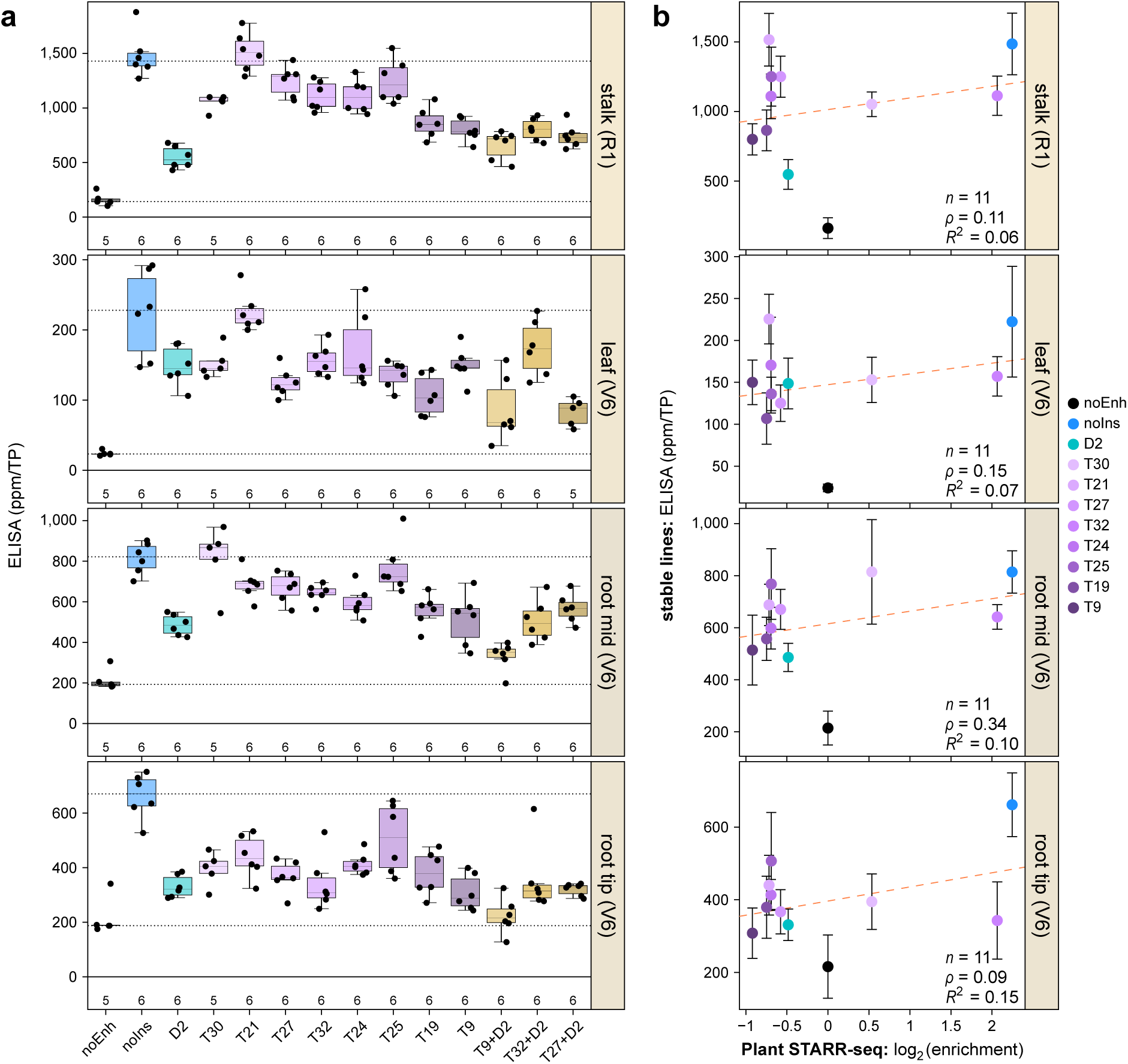
Activity of insulator fragment combinations in different maize tissues. **a**,**b**, Transgenic maize lines were created using insulator fragment combinations in constructs as in Fig. 2h. The activity of insulator fragments was measured in the indicated tissues (**a**) and compared to the corresponding results from Plant STARR-seq in maize protoplasts (**b**). Box plots in **a** are as defined in Fig. 2.

**Extended Data Fig. 5.**
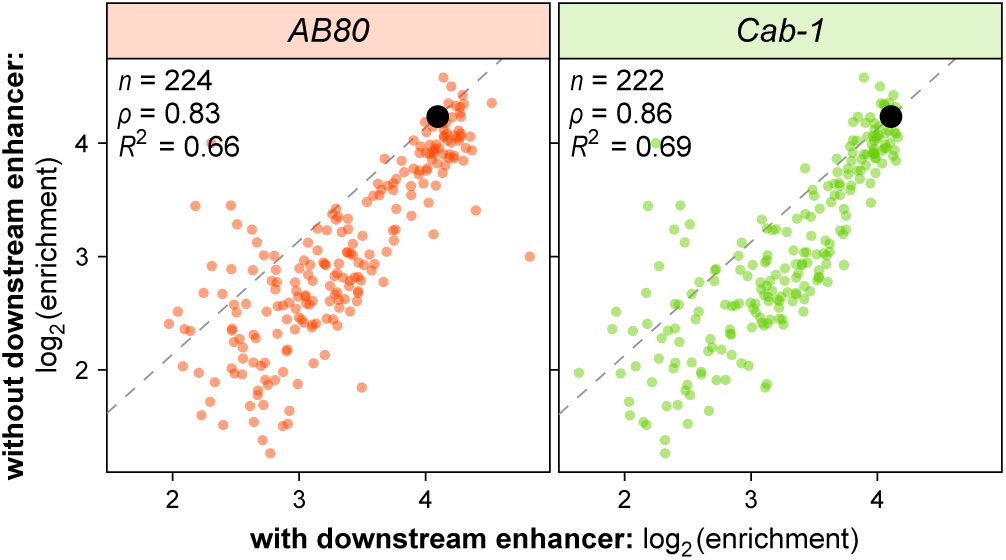
Enhancers downstream of insulator fragments slightly reduce their activity. Correlation between the activity of insulator fragments cloned between a 35S enhancer and a 35S minimal promoter with or without an additional *AB80* or *Cab-1* enhancer inserted between the insulator fragment and 35S minimal promoter. The dashed line represents a y = x line fitted through the point corresponding to a control construct without an insulator (black dot). Pearson’s *R*^2^, Spearman’s *ρ*, and number (*n*) of constructs are indicated.

## References

1. Weising K., Kahl G. Towards an Understanding of Plant Gene Regulation: The Action of Nuclear Factors. Z. Für Naturforschung C 46, 1–11 (1991).

2. Ricci W.A. et al. Widespread long-range cis-regulatory elements in the maize genome. Nat. Plants 5, 1237–1249 (2019).

3. Jores T. et al. Identification of Plant Enhancers and Their Constituent Elements by STARR-seq in Tobacco Leaves. Plant Cell 32, 2120–2131 (2020).

4. Jores T. et al. Synthetic promoter designs enabled by a comprehensive analysis of plant core promoters. Nat. Plants 7, 842–855 (2021).

5. Cuperus J.T. Single-cell genomics in plants: current state, future directions, and hurdles to overcome. Plant Physiol. 188, 749–755 (2022).

6. Schmitz R.J., Grotewold E., Stam M. Cis-regulatory sequences in plants: Their importance, discovery, and future challenges. Plant Cell 34, 718–741 (2022).

7. Jores T., Hamm M., Cuperus J.T., Queitsch C. Frontiers and techniques in plant gene regulation. Curr. Opin. Plant Biol. 75, 102403 (2023).

8. Chetverina D., Aoki T., Erokhin M., Georgiev P., Schedl P. Making connections: Insulators organize eukaryotic chromosomes into independent cis-regulatory networks. BioEssays 36, 163–172 (2014).

9. Heger P., Wiehe T. New tools in the box: An evolutionary synopsis of chromatin insulators. Trends Genet. 30, 161–171 (2014).

10. Burgess-Beusse B. et al. The insulation of genes from external enhancers and silencing chromatin. Proc. Natl. Acad. Sci. 99, 16433–16437 (2002).

11. Vogelmann J., Valeri A., Guillou E., Cuvier O., Nollmann M. Roles of chromatin insulator proteins in higher-order chromatin organization and transcription regulation. Nucleus 2, 358–369 (2011).

12. Singer S.D., Liu Z., Cox K.D. Minimizing the unpredictability of transgene expression in plants: the role of genetic insulators. Plant Cell Rep. 31, 13–25 (2012).

13. Kurbidaeva A., Purugganan M. Insulators in Plants: Progress and Open Questions. Genes 12, 1422 (2021).

14. Hily J.-M., Singer S.D., Yang Y., Liu Z. A transformation booster sequence (TBS) from Petunia hybrida functions as an enhancer-blocking insulator in Arabidopsis thaliana. Plant Cell Rep. 28, 1095–1104 (2009).

15. van der Geest A.H.M., Hall T.C. The β-phaseolin 5′ matrix attachment region acts as an enhancer facilitator. Plant Mol. Biol. 33, 553–557 (1997).

16. Singer S.D., Cox K.D. A gypsy-like sequence from Arabidopsis thaliana exhibits enhancer-blocking activity in transgenic plants. J. Plant Biochem. Biotechnol. 22, 35–42 (2013).

17. Gudynaite-Savitch L., Johnson D.A., Miki B.L.A. Strategies to mitigate transgene–promoter interactions. Plant Biotechnol. J. 7, 472–485 (2009).

18. Singer S.D., Hily J.-M., Liu Z. A 1-kb Bacteriophage Lambda Fragment Functions as an Insulator to Effectively Block Enhancer–Promoter Interactions in Arabidopsis thaliana. Plant Mol. Biol. Report. 28, 69 (2009).

19. Ogbourne S., Antalis T.M. Transcriptional control and the role of silencers in transcriptional regulation in eukaryotes. Biochem. J. 331, 1–14 (1998).

20. Laimins L., Holmgren-König M., Khoury G. Transcriptional “silencer” element in rat repetitive sequences associated with the rat insulin 1 gene locus. Proc. Natl. Acad. Sci. 83, 3151–3155 (1986).

21. Pang B., Snyder M.P. Systematic identification of silencers in human cells. Nat. Genet. 52, 254–263 (2020).

22. Gisselbrecht S.S. et al. Transcriptional Silencers in *Drosophila* Serve a Dual Role as Transcriptional Enhancers in Alternate Cellular Contexts. Mol. Cell 77, 324–337.e8 (2020).

23. Jores T. et al. Plant enhancers exhibit both cooperative and additive interactions among their functional elements. Plant Cell 36, 2570–2586 (2024).

24. Gorjifard S., et al. Arabidopsis and Maize Terminator Strength is Determined by GC Content, Polyadenylation Motifs and Cleavage Probability. Preprint at https://www.biorxiv.org/content/10.1101/2023.06.16.545379v2 (2024).

25. She W. et al. The gypsy Insulator of Drosophila melanogaster, Together With Its Binding Protein Suppressor of Hairy-Wing, Facilitate High and Precise Expression of Transgenes in Arabidopsis thaliana. Genetics 185, 1141–1150 (2010).

26. Gdula D.A., Gerasimova T.I., Corces V.G. Genetic and molecular analysis of the gypsy chromatin insulator of Drosophila. Proc. Natl. Acad. Sci. 93, 9378–9383 (1996).

27. Parkhurst S.M. et al. The Drosophila su(Hw) gene, which controls the phenotypic effect of the gypsy transposable element, encodes a putative DNA-binding protein. Genes Dev. 2, 1205–1215 (1988).

28. Zhao K., Hart C.M., Laemmli U.K. Visualization of chromosomal domains with boundary element-associated factor BEAF-32. Cell 81, 879–889 (1995).

29. Gaszner M., Vazquez J., Schedl P. The Zw5 protein, a component of the scs chromatin domain boundary, is able to block enhancer–promoter interaction. Genes Dev. 13, 2098–2107 (1999).

30. Bell A.C., West A.G., Felsenfeld G. The Protein CTCF Is Required for the Enhancer Blocking Activity of Vertebrate Insulators. Cell 98, 387–396 (1999).

31. Banerji J., Rusconi S., Schaffner W. Expression of a β-globin gene is enhanced by remote SV40 DNA sequences. Cell 27, 299–308 (1981).

32. Antes T.J., Namciu S.J., Fournier R.E.K., Levy-Wilson B. The 5‘ Boundary of the Human Apolipoprotein B Chromatin Domain in Intestinal Cells. Biochemistry 40, 6731–6742 (2001).

33. West A.G., Gaszner M., Felsenfeld G. Insulators: many functions, many mechanisms. Genes Dev. 16, 271–288 (2002).

34. Singer S.D., Hily J.-M., Cox K.D. Analysis of the enhancer-blocking function of the TBS element from Petunia hybrida in transgenic Arabidopsis thaliana and Nicotiana tabacum. Plant Cell Rep. 30, 2013–2025 (2011).

35. Halperin S.O. et al. CRISPR-guided DNA polymerases enable diversification of all nucleotides in a tunable window. Nature 560, 248–252 (2018).

36. Gibson D.G. et al. Enzymatic assembly of DNA molecules up to several hundred kilobases. Nat. Methods 6, 343–345 (2009).

37. Engler C., Kandzia R., Marillonnet S. A One Pot, One Step, Precision Cloning Method with High Throughput Capability. PLOS ONE 3, e3647 (2008).

38. Debernardi J.M. et al. A GRF–GIF chimeric protein improves the regeneration efficiency of transgenic plants. Nat. Biotechnol. 38, 1274–1279 (2020).

39. Tonnies J., Mueth N.A., Gorjifard S., Chu J., Queitsch C. Scalable Transfection of Maize Mesophyll Protoplasts. JoVE J. Vis. Exp. e64991 (2023).

40. Anand A. et al. High efficiency Agrobacterium-mediated site-specific gene integration in maize utilizing the FLP-FRT recombination system. Plant Biotechnol. J. 17, 1636–1645 (2019).

41. Clough S.J., Bent A.F. Floral dip: a simplified method for Agrobacterium -mediated transformation of Arabidopsis thaliana. Plant J. 16, 735–743 (1998).

42. Hiei Y., Komari T. Agrobacterium-mediated transformation of rice using immature embryos or calli induced from mature seed. Nat. Protoc. 3, 824–834 (2008).

43. Masella A.P., Bartram A.K., Truszkowski J.M., Brown D.G., Neufeld J.D. PANDAseq: paired-end assembler for illumina sequences. BMC Bioinformatics 13, 1–7 (2012).

