## Supplementary Table 1 for "Small DNA elements that act as both insulators and silencers in plants"

**Supplementary Table 1 | Insulator fragments used in fragment combination library.**

| **insulator** | **start** | **stop** | **orientation** | **insulator activity in tobacco** |
| --- | --- | --- | --- | --- |
| **fragments for positions 1 and 2** | | | | |
| β-phaseolin | 230 | 399 | fwd and rev | bottom 25% (both orientations) |
| β-phaseolin | 383 | 552 | fwd and rev | bottom 25% (both orientations) |
| β-phaseolin | 1148 | 1317 | fwd and rev | top 25% (rev orientation) |
| β-phaseolin | 1317 | 1486 | fwd and rev | top 25% (rev orientation) |
| β-phaseolin | 1395 | 1564 | fwd and rev | top 25% (both orienations) |
| β-phaseolin | 1633 | 1802 | fwd and rev | top 25% (fwd orientation) |
| β-phaseolin | 1712 | 1881 | fwd and rev | top 25% (both orienations) |
| β-phaseolin | 1791 | 1960 | fwd and rev | top 25% (both orienations) |
| β-phaseolin | 2266 | 2435 | fwd and rev | top 25% (both orienations) |
| β-phaseolin | 2345 | 2514 | fwd and rev | top 25% (fwd orientation) |
| β-phaseolin | 2741 | 2910 | fwd and rev | top 25% (fwd orientation) |
| β-phaseolin | 3058 | 3227 | fwd and rev | top 25% (both orienations) |
| β-phaseolin | 3454 | 3623 | fwd and rev | bottom 25% (both orientations) |
| TBS | 252 | 421 | fwd and rev | top 25% (both orienations) |
| TBS | 588 | 757 | fwd and rev | top 25% (both orienations) |
| TBS | 756 | 925 | fwd and rev | bottom 25% (both orientations) |
| TBS | 1681 | 1850 | fwd and rev | top 25% (both orienations) |
| TBS | 1765 | 1934 | fwd and rev | top 25% (rev orientation) |
| λ-EXOB | 1 | 170 | fwd and rev | top 25% (both orienations) |
| λ-EXOB | 83 | 252 | fwd and rev | top 25% (both orienations) |
| λ-EXOB | 166 | 335 | fwd and rev | top 25% (both orienations) |
| λ-EXOB | 249 | 418 | fwd and rev | top 25% (both orienations) |
| λ-EXOB | 332 | 501 | fwd and rev | top 25% (both orienations) |
| λ-EXOB | 415 | 584 | fwd and rev | top 25% (both orienations) |
| λ-EXOB | 663 | 832 | fwd and rev | top 25% (rev orientation) |
| λ-EXOB | 829 | 998 | fwd and rev | top 25% (fwd orientation) |
| BEAD-1C | 246 | 415 | fwd and rev | top 25% (both orienations) |
| UASrpg | 157 | 326 | fwd and rev | top 25% (fwd orientation) |
| sIns1 | 54 | 223 | fwd and rev | top 25% (rev orientation) |
| sIns2 | 335 | 504 | fwd and rev | top 25% (both orienations) |
| gypsy | 1 | 170 | fwd and rev | bottom 25% (both orientations) |
| gypsy | 54 | 223 | fwd and rev | bottom 25% (both orientations) |
| **fragments for position 3** | | | | |
| β-phaseolin | 1633 | 1802 | fwd | top 5% |
| β-phaseolin | 1712 | 1881 | rev | top 5% |
| λ-EXOB | 1 | 170 | fwd | top 5% |
| λ-EXOB | 332 | 501 | fwd | top 5% |
| λ-EXOB | 415 | 584 | rev | top 5% |
