## Supplementary Table 2 for "Small DNA elements that act as both insulators and silencers in plants"

**Supplementary Table 2 | Insulator fragment combinations tested in stable transgenic maize lines.** Fragments are numbered by increasing distance from the minimal promoter (fragment 1 is the fragment closest to the promoter, fragment 3 the most distal one)

| **name** | **number of fragments** | **fragment 3** | **fragment 2** | **fragment 1** | **insulator activity in maize protoplasts** |
| --- | --- | --- | --- | --- | --- |
| **D2** | 2 |  | β-phaseolin 1148-1317, fwd | sIns2 335-504, fwd | strong |
| **T30** | 3 | β-phaseolin 1633-1802, fwd | λ-EXOB 663-832, fwd | β-phaseolin 1564-1395, rev | intermediate |
| **T21** | 3 | β-phaseolin 1881-1712, rev | λ-EXOB 663-832, fwd | λ-EXOB 170-1, rev | weak |
| **T27** | 3 | λ-EXOB 1-170, fwd | λ-EXOB 584-415, rev | TBS 925-756, rev | strong |
| **T32** | 3 | λ-EXOB 584-415, rev | λ-EXOB 832-663, rev | β-phaseolin 1960-1791, rev | strong |
| **T24** | 3 | λ-EXOB 584-415, rev | β-phaseolin 3227-3058, rev | λ-EXOB 170-1, rev | strong |
| **T25** | 3 | λ-EXOB 1-170, fwd | β-phaseolin 1802-1633, rev | β-phaseolin 1317-1148, rev | strong |
| **T19** | 3 | λ-EXOB 1-170, fwd | β-phaseolin 1148-1317, fwd | sIns2 335-504, fwd | strong |
| **T9** | 3 | λ-EXOB 584-415, rev | TBS 1765-1934, fwd | sIns2 335-504, fwd | strong |
