## Supplementary Table 3 for "Small DNA elements that act as both insulators and silencers in plants"

**Supplementary Table 3 | Insulators and insulator fragments used in the enhancer-insulator combination library.**

| **type** | **insulator** | **start** | **stop** | **orientation** |
| --- | --- | --- | --- | --- |
| full-length insulator | λ-EXOB | 1 | 998 | fwd |
| full-length insulator | BEAD-1C | 1 | 538 | fwd |
| full-length insulator | UASrpg | 1 | 378 | fwd |
| full-length insulator | sIns1 | 1 | 386 | fwd |
| full-length insulator | sIns2 | 1 | 504 | fwd |
| full-length insulator | gypsy | 1 | 386 | fwd |
| insulator fragment | β-phaseolin | 1395 | 1564 | fwd |
| insulator fragment | TBS | 756 | 925 | fwd |
| insulator fragment | λ-EXOB | 1 | 170 | fwd |
| insulator fragment | BEAD-1C | 246 | 415 | fwd |
| insulator fragment | UASrpg | 1 | 170 | fwd |
| insulator fragment | gypsy | 54 | 223 | fwd |
