## Supplementary Table 4 for "Small DNA elements that act as both insulators and silencers in plants"

**Supplementary Table 4 | Full-length insulator sequences.**

| **insulator** | **sequence** |
| --- | --- |
| β-phaseolin | CATAAGAAATATGAAAATCGTTATGAACTTTATTATTTGTTAAACGTTTTCATAACCGCATAAAA  TTTTATAAAGTCCCGTCTATCTTTAATATGTAGTCTAACATTTTCATATTGAAATATATAATTTA  CTTAATTTTAGTGTTGGTAGAAAGCATAATGATTTATTCTTGTTCATATAAATGTTTAATATAAA  CAAACTCTTTACCTTAAGAAGGATTTCCCATTTTATATTTTAAAAATATATTTATCAAATATTTT  TCAACCACGTAAATCTCATAATAATAAGTTGTTTCAAAAGTAATAAAATTTAACTCCATAATTTT  TTTATTTGACTGATCTTAAAGCAACACCCAGTGACACAACTAGTAATTTTTTTCTTTGAATAAAA  AAATCCAATTATCATTGTATTTTTTTTATACAATGAAAATTTCACCAAACAATGATTTGTGGTAT  TTCTGAAGCAAGTCATGTTAATGCAAAATTCTATAATTCACATTTGACACTACGGAAGTGACTGA  AAATCTGTTTTTACATGCGAGACACATCAATTTTTAATTCTAAAGTAATTTTAATAATAGTTACT  ATATTCAAGATTTGATATATCAAATACTCAATATTACTTCTAAAAAATTAATTAGATATAATTAA  AAAATTACTTTTTTAATTTTAAGTTTAATTGTTGGATTTGTGACTATTGATTTATTATTCTACTA  TGTTTAAACTGTTTTATAGATAGTTTAAAGTAAATATAAGTATTGTAGAGTGTTACCGTAAACTA  TAAGATTTATGTTGGACTAATTTTATGTTCTTCATTTGCAATATTTTAATATATTTGTTGTTGGT  TTACCTTTCTTGGTATGTAAGTCCGTAACCAGAATTACTGTGGGTTGCCATGGCACTCTGTAGTC  TTTTGGTTCGTGCATGGATGCTTGCGCAAGAAAAAGACATAGAACAAAGAAAAAAGACAAAACAG  AGAGAGAAAACGCAATCACACAACCAACTCAAATTAGTCACTGGCTGATCAAGATCGCCGCGTCC  ATGTATGTCTAAATGCCATGCAAAGCAACACGTGCTTAACATGCACTTTAAATGGCTCACCCATC  TCAACCCACACACAAACACATTGTCTTTTTCTTCATCATCACCACAACCACCTGTATATATTCAT  TCTCTTCCGCCACCTCAATTTCTTCACTTCAACACACGTCAACCTGCATATGCGTGTCATCCCAT  GCCCAAATCTCCATGCATGTTCCAACCACCTTCTCTCTTATATAATACCTATAAATACCCCTAAT  ATCACTCACTTCTTTCATCATCCATCCATCCAGAGTACTACTACTCTACTACTATAATACCCCAA  CCCAACTCATATTCAATACTACTCTACTATGATGAGAGCAAGGGTTCCACTCCTGTTGCTGGGAA  TTCTTTTCCTGGCATCACTTTCTGCCTCATTTGCCACTTCACTCCGGGAGGAGGAAGAGAGCCAA  GATAACCCCTTCTACTTCAACTCTGACAACTCCTGGAACACTCTATTCAAAAACCAATATGGTCA  CATTCGTGTCCTCCAGAGGTTCGACCAACAATCCAAACGACTTCAGAATCTTGAAGACTACCGTC  TTGTGGAGTTCAGGTCCAAACCCGAAACCCTCCTTCTTCCTCAGCAGGCTGATGCTGAGTTACTC  CTAGTTGTCCGTAGTGGTAAGTAATTGCTACTGGTATCACTTGTTTCTTCTTGCAGAAATAATGG  TAATGAGTTTTTTTATAATTTCAGGGAGCGCCATACTCGTCTTGGTGAAACCTGATGATCGCAGA  GAGTACTTCTTCCTTACGCAAGGCGATAACCCGATATTCTCTGATAACCAGAAAATCCCTGCAGG  AACCATTTTCTATTTGGTTAACCCTGACCCCAAAGAGGATCTCAGAATAATCCAACTCGCCATGC  CCGTTAACAACCCTCAGATTCATGTACTGCCTTTTGTAATACCAAACTAATTTTTTTGTTATTTT  AACTTGCAATTTCTCTCCAAATGTGATGATAAATGTTTGTCCTGTAGGAATTTTTCCTATCTAGC  ACAGAAGCCCAACAATCCTACTTGCAAGAGTTCAGCAAGCATATTCTAGAGGCCTCCTTCAATGT  AAGAAAGAAAACAGCATCTAACTACATATTTGCGTCATCTAACTACATATTTTCGTTGCCATTTA  GCTAGTACTTTGTCTAAATGTCACACTTGTTGAATTTGTTGAATGATATCATTATATATGTTTGC  ATGATTTTTATAGAGCAAATTCGAGGAGATCAACAGGGTTCTGTTTGAAGAGGAGGGACAGCAAG  AGGGAGTGATTGTGAACATTGATTCTGAACAGATTGAGGAACTGAGCAAACATGCAAAATCTAGT  TCAAGGAAATCCCATTCCAAACAAGATAACACAATTGGAAACGAATTTGGAAACCTGACTGAGAG  GACCGATAACTCCTTGAATGTGTTAATCAGTTCTATAGAGATGAAAGAGGTAAATACAAAGAAAA  AACATATAGACAAACTTAGCAATTGAGTTCTATTATTCACTGTCGTCTTGGTTAGAAAATCTTAG  TATTGAGAATATAATTAAATAATGGTTTTTTTTGTTAACAAATTTAGGGAGCTCTTTTTGTGCCA  CACTACTATTCTAAGGCCATTGTTATACTAGTGGTTAATGAAGGAGAAGCACATGTTGAACTTGT  TGGCCCAAAAGGAAATAAGGAAACCTTGGAATTTGAGAGCTACAGAGCTGAGCTTTCTAAAGACG  ATGTATTTGTAATCCCAGCAGCATATCCAGTTGCCATCAAGGCTACCTCCAACGTGAATTTCACT  GGTTTCGGTATCAATGCTAATAACAACAATAGGAACCTCCTTGCAGGTATATATATTTATTATAT  ATGACCATGAATTTGAATATAGGGTTGTTGATGGGATTTTTTATTTATAATTGGTAATGCGTGAT  TGTGATTGAAAATATGAAGGTAAGACGGACAATGTCATAAGCAGCATCGGTAGAGCTCTGGACGG  TAAAGACGTGTTGGGGCTTACGTTCTCTGGGTCTGGTGAAGAAGTTATGAAGCTGATCAACAAGC  AGAGTGGATCGTACTTTGTGGATGGACACCATCACCAACAGGAACAGCAAAAGGGAAGTCACCAA  CAGGAACAGCAAAAGGGAAGAAAGGGTGCATTTGTGTACTGAATAAGTATGAACTAAAATGCATG  TATGGTGTAAGAGCTCATGGAGAGCATGGAAATATGTATCAGACCATGTAACACTATAATAACTG  AGCTCCATCTCACTTCTTCTATGAATAAACAAAGGATGTTATGATATATTAACACTATATGCACC  TTACATAGTAATACATTAATATTTAATACTTTTTATTTTAACTTTTTAGTTTAAAATATTATTAT  ATTATTAACTTTTTAGTTTAAAATATTTATATTATTATAAAGAGAAATAAACAAAGGATGTTATG  ATATATTAACACTATATGTACCTTACATAGTAATATATTAATATTTAATACTTTTTATTTTAACT  TTTTAATTTAAAATATTATTATAAATGATGCTTGTGTTTTATGTGTTGGCATGCTTGTATTTTAT  GTGTTGACTTTCTGTGTGAAGGTAATGTGATATGGTTAGCTGGTGGTAACAATTGTGTTTTATGT  GTTGGCTTTCTGTGAAGCTAATTTGATATGGTTAGCTGATGGGAACAAAATATTAAAGGAAGCTA  ATTTGATATGGTTAGCTGATAGTAACAAAATATCAAAATAAATTTCTTCTTACTTTAATAAATTA  TATGAATTGTGACGGATTATATGGAATGTATAGGACAAAATCTTTAATAAATTACATGAATTGTG  ACGGATTATGGAATGGAATGTAGCAAATAGGACAAAACAAATGTTTGTAAGAACCAAGAGATCCT  AACCATGTATAGGCTAACCATATATAGGCTTAGGCCAAAAACAAACCTAAACTCTTAAACTTTGA  TTTATTATAAAAAAAAAATGATTATTTAATATATAGGCTTAGGCCAAAAACAAACCTAAACTCTT  AAACTTTGATTTATTATAAAAAAAAAATGATTATTTAACACCATGCACCTTACACCCTACTAAAT  TTTAATATTTTAATTATTTTTAATTTTAAAAATTATTTTAATATTTTACATTATTTTCACTAAAA  ACATTTGTTTTTTTTATTATAAATATTAATATTTTATTATAGATATAAAGTGAAGCAATATTGTT  TGAAAATATCATTACCCGTTACAAAATATTGTACAACTAGCTATAAAAAAGCAAACCACAAGGAA  ACAGAAGACTTTCACTTTGAAAAGGGGTGCCTGCTAAGACCGTAATTTTGCTTCAGATAAAAAAC  CATCACAACTCAATAGGTACTCACAAAACTACTT |
| TBS | TTCCTAACACCTGGAGAACCTTTTATGTACTTCACAACCCTCAATGCTGCTTCCAGGTGTGACTG  TTTGGGTTGCTGCATGAATTGACTCAGCACCTGCACTGCAAAGCTGATATCTGGCCTTGTGATTG  TCAAGTAGAGAAGCTTTCCCACCAGTCTCTGATAGGATCCCACATCCTCTAACTCTGCATCATCA  GTCTTTCCTAAGTGCTTGTCATATTCTACAGTAGTCAGCCTTTGGTTCTGTTCCATTGGGGTTGA  CACTGGTCTGCAGCCTCCCAGACCAACACCTGATATCAGTTCCAGTGCATACTTCCTCTGGTTCA  GTAGGATTCCCTTTTCTGATCTCAGCACCTCAATGCCTAAGAAGTATTTTAGTTCTCCCAAATCT  TTCATCTTGAAATGCTGATGCAGGGTTGCCTTTGCTTCTGAAATCAAAACATTGCTGCTGCCTGT  TATTAACAGATCATCCACATAAATCAGGATTATGACAAGGTCAGTCCCCTCTCTTTTGGTAAACA  AGGAGTGATCATAAGCACTTTGCATAAAACCAGCCTGCATAAGGACAGTGGTAAGCTTGATGTTC  CACTGCCTTGATGCTTGCTTTAAACCATAGAGGGATTCAACAGCCTGCACACTTTGGACTCCCCT  TGGCTGTGAAAACCCTGAGGCAGAGACATATAAACTTCTTCCATGAGGTCACCTTGTAGAAAAGC  ATTGTTGACATCCATCTGGAAAAGGAACCAGCCCTTGGAAGCAGCAACAGATATGACAGCTTTTA  CAGTGACCATTTTGGCCACTGGAGAAAAAGTTTCATGGTAGTCAAGGCCTTCTTGCTGAGTGTAT  CCCTTGGCCACTAGCCTTGCCTTAAACCTGTCAACTTCACCATTAGCTTTGTATTTAATTTTGTA  CACCCATTTGGACCCTATAGGCTGTTTACCAGGGGGTAAAGGGACAATCTCCCAGGTGTTATTAT  CCTCAAGAGCCTGTATCTCAAGGGACATGGCCTCCATCCATTTCTCATCTTGAGCTGCTTCTTTG  AAGGATTTAGGTTCAGTATAAGTGGAGAAAGCACTCAAATAAGCTTGATAGTGAACAGGCAAGTG  ATCATAGGAAACATAGTTGGCTATAGGGTATGGAACATCCCTAGAGCCTTTGTTCAGTGTCACAA  AGTCCTTGAGCCAGATGGGAGGACCTGCATTGCGTTTAGGTCTAGTGTGCAGGTTTGCTGGAACT  AGAGAGGGATCAGCTACAGCAGTATGGTGCTCAAATTCAGCATTTGCTAGGTCAGGCTCAGCTGA  CTCAACTGACTCTGCAGGAGCATGCAGGTGGCCTGAAGGTGCAGCATCAGCTGAAGTGATATGAT  TTTGGGTAGAATGAGTCCCCAAAGTGATATCCTCATTGGCCTCTACAATGTCAGGAATAAGAACT  GGAGTATCATCATCATCAGTATGAAAGCTGGAAGGAAAAATATCATTATACACTAGCTGCAAAGC  TGTATCTTCAGAGCTAAGTGAACCTGCTCGAGTGAACATATCAGGTTCATGGGAGATGGACTCTT  TGAAAGGGAACTGAAACTCTCTGAAAACTACATCCCTGCTCACTATGATCACCTTATTATCCAAG  TCATACAACCTGTAACCCTTTTGAGTCTCAGAATAACCAATGAAGATGGTTCGCTTGGCTCTAGG  TGCAAGTTTGTCACCTTTGGGCAGTGTTGCTGCAAAAGCAAGACACCCAAACACTCTCAAGTGAT  CAAGCTTGGCATGTTTCTGATAGAGTAGTTCATATGGACATTTGCCTTGTAAGATTGGAGTAGGG  AGTCTATTGATCAGGTATACAACAGTCTTGACACAGTCTCCCCAAAACCTGGTAGGTACACCACT  CTGAAACTTAAGTGCCCTTGCCATCTCAAGGATGTGTCTGTGCTTTCTCTCCACAACACCATTCT  GTTGTGGTGTGTAGGGACAGCTACTTTGATGAACAATCCCAAGAGAGGCCAGCAACTCATTACAA  CTT |
| λ-EXOB | AATTCAAACAGGGTTCTGGCGTCGTTCTCGTACTGTTTTCCCCAGGCCAGTGCTTTAGCGTTAAC  TTCCGGAGCCACACCGGTGCAAACCTCAGCAAGCAGGGTGTGGAAGTAGGACATTTTCATGTCAG  GCCACTTCTTTCCGGAGCGGGGTTTTGCTATCACGTTGTGAACTTCTGAAGCGGTGATGACGCCG  AGCCGTAATTTGTGCCACGCATCATCCCCCTGTTCGACAGCTCTCACATCGATCCCGGTACGCTG  CAGGATAATGTCCGGTGTCATGCTGCCACCTTCTGCTCTGCGGCTTTCTGTTTCAGGAATCCAAG  AGCTTTTACTGCTTCGGCCTGTGTCAGTTCTGACGATGCACGAATGTCGCGGCGAAATATCTGGG  AACAGAGCGGCAATAAGTCGTCATCCCATGTTTTATCCAGGGCGATCAGCAGAGTGTTAATCTCC  TGCATGGTTTCATCGTTAACCGGAGTGATGTCGCGTTCCGGCTGACGTTCTGCAGTGTATGCAGT  ATTTTCGACAATGCGCTCGGCTTCATCCTTGTCATAGATACCAGCAAATCCGAAGGCCAGACGGG  CACACTGAATCATGGCTTTATGACGTAACATCCGTTTGGGATGCGACTGCCACGGCCCCGTGATT  TCTCTGCCTTCGCGAGTTTTGAATGGTTCGCGGCGGCATTCATCCATCCATTCGGTAACGCAGAT  CGGATGATTACGGTCCTTGCGGTAAATCCGGCATGTACAGGATTCATTGTCCTGCTCAAAGTCCA  TGCCATCAAACTGCTGGTTTTCATTGATGATGCGGGACCAGCCATCAACGCCCACCACCGGAACG  ATGCCATTCTGCTTATCAGGAAAGGCGTAAATTTCTTTCGTCCACGGATTAAGGCCGTACTGGTT  GGCAACGATCAGTAATGCGATGAACTGCGCATCGCTGGCATCACCTTTAAATGCCGTCTGGCGAA  GAGTGGTGATCAGTTCCTGTGGG |
| BEAD-1C | TTCAGTAATACGGGTAGCTGGGACATGCCATATTTGGAACACATTTATACTAAAAAAGTATTCAT  TGTTTATCTGAAATTCAAATTCCACTGGGCATCCTGTGTTTTATCTGGCAATGCTAGGCATGCAG  AATACCAAAAGTAAGCACCAGGCAGGCCAGAGTCCCACCATGAGCATCTTCAGGGCCCCTGGATG  TGGAAGAGGGATGTTGAGGGCCCAGGGGCTGCCTTGCCGGTGCATTGGCTGCCCAGGCCTGCACT  GCCGCCTGCCGGCAGGGGTCCAGTCCACGAGTCCCAGCTCCCTGCTGGCGGAAGTCCATTTCAGA  GCTTCCGGTTCTCCCAAGTCCAAGGATTATGCTCACTCCCCACCCACAGTCTCTTAGTGTCTGTC  CCTGCTCTAAAGATGTTGTCTGGGCTTGATATTAATATGAGAGCTGACTGTTCCCTTCCTGATCT  AGACCATAACCATCTTCAAGTTAAATTGCTCCTCCTCTTCTAACTGCCCAACCTCACCCACGTCT  GACCATACCCAAGCACAG |
| UASrpg | AGCTTGCCTCGTCCCGCCGGGTCACCCGGCCAGCGACATGGAGGCCCAGAATACCCTCCTTGACA  GTCTTGACGTGCGCAGCTCAGGGGCATGATGTGACTGTCGCCCGTACATTTAGCCCATACATCCC  CATGTATAATCATTTGCATCCATACATTTTGATGGCCGCACGGCGCGAACGAAAAATTACGGCTC  CTCGCTGCAGACCTGCGAGCAGGGAAACGCTCCCCTCACAGACGCGTTGAATTCTCCCCACGGCG  CGCCCCTGTAGAGAAATATAAAAGGTTAGGATTTGCCACTGAGGTTCTTCTTTCATATACTTCCT  TTTAAAATCTTGCTAGGATACAGTTCTCACATCACATCCGAACATAAACAAAA |
| gypsy | CACGTAATAAGTGTGCGTTGAATTTATTCGCAAAAACATTGCATATTTTCGGCAAAGTAAAATTT  TGTTGCATACCTTATCAAAAAATAAGTGCTGCATACTTTTTAGAGAAACCAAATAATTTTTTATT  GCATACCCGTTTTTAATAAAATACATTGCATACCCTCTTTTAATAAAAAATATTGCATACTTTGA  CGAAACAAATTTTCGTTGCATACCCAATAAAAGATTATTATATTGCATACCCGTTTTTAATAAAA  TACATTGCATACCCTCTTTTAATAAAAAATATTGCATACGTTGACGAAACAAATTTTCGTTGCAT  ACCCAATAAAAGATTATTATATTGCATACCTTTTCTTGCCATACCATTTAGCCGATCAATT |
| sIns1 | AGAACTTTAAAAGTGCTCATCATTGGAAAACGTTCTTCGGGGCGAAAACTCTCAAGGATCTTACC  GCTGTTGAGATCCAGTTCGATGTAACCCACTCGTGCACCCAACTGATCTTCAGCATCTTTTACTT  TCACCAGCGTTTCTGGGTGAGCAAAAACAGGAAGGCAAAATGCCGCAAAAAAGGGAATAAGGGCG  ACACGGAAATGTTGAATACTCATACTCTTCCTTTTTCAATATTATTGAAGCATTTATCAGGGTTA  TTGTCTCATGAGCGGATACATATTTGAATGTATTTAGAAAAATAAACAAATAGGGGTTCCGCGCA  CATTTCCCCGAAAAGTGCCACCTGACGTCCAACATATGGCACCGGAGGCTTTCGTCTTCAC |
| sIns2 | GCTGAACGCAAAGCTGATCACTCAGCGGAAGTTCGACAATCTCACTAAGGCTGAGAGGGGCGGAC  TGAGCGAACTGGACAAAGCAGGATTCATTAAACGGCAACTTGTGGAGACTCGGCAGATTACTAAA  CATGTCGCCCAAATCCTTGACTCACGCATGAATACCAAGTACGACGAAAACGACAAACTTATCCG  CGAGGTGAAGGTGATTACCCTGAAGTCCAAGCTGGTCAGCGATTTCAGAAAGGACTTTCAATTCT  ACAAAGTGCGGGAGATCAATAACTATCATCATGCTCATGACGCATATCTGAATGCCGTGGTGGGA  ACCGCCCTGATCAAGAAGTACCCAGCACTGGAAAGCGAGTTCGTGTACGGAGACTACAAGGTCTA  CGACGTGCGCAAGATGATTGCCAAATCTGAGCAGGAGATCGGAAAGGCCACCGCAAAGTACTTCT  TCTACAGCAACATCATGAATTTCTTCAAGACCGAAATCACCCTTGCAAA |
